# Autophosphorylation of the oncogenic protein TEL-ABL confers resistance to the allosteric ABL inhibitor asciminib

**DOI:** 10.1101/2024.10.02.616342

**Authors:** Serena Muratcioglu, Christopher A. Eide, Kent Gorday, Emily Sumpena, Wenqi Zuo, Jay T. Groves, Brian J. Druker, John Kuriyan

**Author notes:** insitro, San Francisco, CA, USA. ^4^Department of Pediatrics, University of California, Los Angeles, Los Angeles, CA, USA.

## Abstract

Chromosomal translocations that fuse the ABL1 gene to BCR and TEL cause human leukemias. Oligomerization and the loss of an inhibitory myristoylation modification lead to unregulated kinase activity of the BCR-ABL and TEL-ABL fusion proteins. ATP-competitive ABL inhibitors, such as imatinib and ponatinib, are effective against both fusion proteins. We discovered that asciminib, an allosteric inhibitor of BCR-ABL that binds to the myristoyl binding site in the ABL kinase domain, is >2000-fold less potent against TEL-ABL than BCR-ABL in cell-growth assays. This is surprising because the ABL components of the two fusion proteins, including the asciminib binding sites, have identical sequence. We deleted a short helical segment in the ABL kinase domain that closes over asciminib when it is bound. This deletion results in asciminib resistance in BCR-ABL, but has no effect on TEL-ABL, suggesting that the native autoinhibitory mechanism that asciminib engages in BCR-ABL is disrupted in TEL-ABL. We show, using mammalian cell expression and single-molecule microscopy, that BCR-ABL is mainly dimeric while TEL-ABL forms higher-order oligomers. Oligomerization can promote trans-autophosphorylation of ABL, and we find that a regulatory phosphorylation site in the SH3 domain of ABL (Tyr 89) is highly phosphorylated in TEL-ABL. This phosphorylation is expected to disassemble the autoinhibited conformation of ABL, thereby preventing asciminib binding. We show that TEL-ABL is intrinsically susceptible to inhibition by asciminib, but that increased phosphorylation results in resistance. Our results demonstrate that different ABL fusion proteins can have dramatically different responses to allosteric inhibitors due to differential phosphorylation.

**One Sentence Summary:** When TEL-ABL is phosphorylated, it is insensitive to asciminib. However, when TEL-ABL is dephosphorylated by a phosphatase, asciminib sensitivity is restored.

## INTRODUCTION

Cancer-causing variants of the cytoplasmic tyrosine kinase c-Abl (ABL1) are generated by fusion of the gene encoding c-Abl with genes for proteins such as breakpoint cluster region (BCR) and translocated ETS leukemia (TEL) (*1*) (Figure 1A). The mechanisms that normally regulate the activity of c-Abl are disrupted in the oncogenic fusion proteins, leading to constitutively active kinases. The best-studied of these fusion proteins is BCR-ABL, which causes chronic myelogenous leukemia (CML) (*2*, *3*), neutrophilic leukemia, and acute lymphocytic leukemia (ALL) (*4*). BCR-ABL accounts for more than 90% of CML cases. Depending on the location of the translocation breakpoint in the BCR gene, distinct variants of the BCR-ABL fusion protein are produced. p210 and p185 BCR-ABL are the most common variants in CML and ALL, respectively (*3*).

**Fig. 1.**
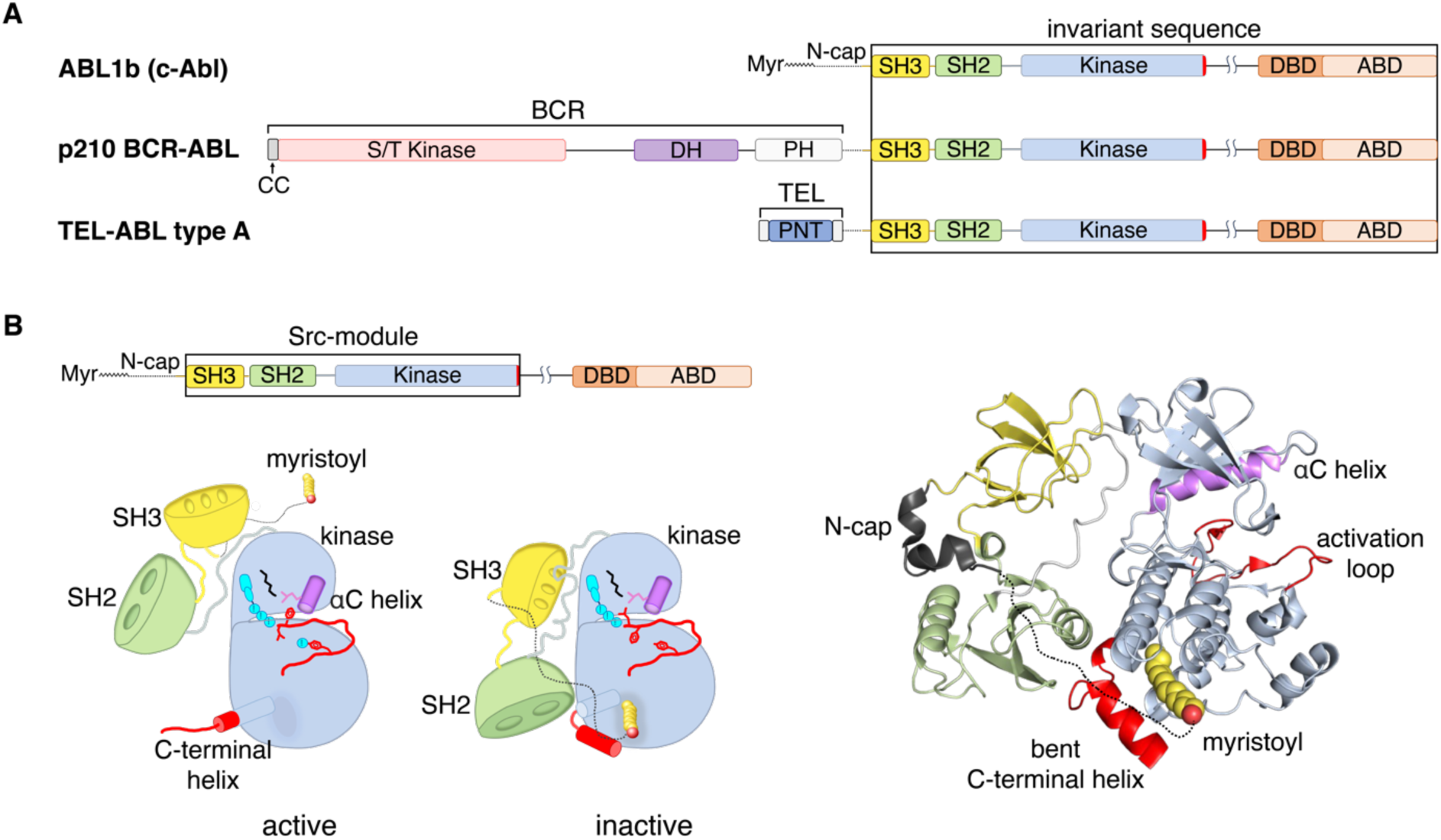
Domain organization of ABL kinase and ABL fusion proteins and conformational states of c-Abl. **(A**) Schematic domain representation of c-Abl, p210 BCR-ABL, and TEL-ABL (type A) fusion proteins. Myr: myristoyl group; N-cap: N-terminal cap; SH2: Src-homology 2, SH3: Src-homology 3.; DBD: DNA-binding domain; ABD: actin-binding domain; CC: coiled-coil domain; S/T kinase: serine/threonine kinase; DH: Dbl-homology; PH: Pleckstrin-homology; PNT: pointed. **(B)** Schematic representation of active and autoinhibited ABL as well as the crystal structure of the autoinhibited ABL “Src-module” in complex with the adenosine triphosphate (ATP)–competitive inhibitor PD166326 (right; PDB entry 2FO0). The structured portion of the myristoylated N-cap is shown as a black cartoon representation. The unstructured portion of the myristoylated N-cap that engages the C-lobe of the kinase domain is shown as a dotted line. Domains in the structure are color-coded and correspond to the schematic shown at the top. In the kinase domain, the positions of helix αC and the activation loop are rendered in magenta and red, respectively. The side chain of the activation loop autophosphorylation site (Tyr 412) is also shown. Myristoyl group is shown in yellow spheres. The distal lobe of the C-terminal helix is shown in red.

TEL (also known as ETS–variant gene 6, ETV6), a member of the ETS (Erythroblast Transformation Specific) family of transcriptional regulators, also forms oncogenic fusion proteins with ABL (*5*–*7*). TEL-ABL fusions are rare in human hematologic malignancies, compared to BCR-ABL fusions. A small number of incidences of TEL-ABL fusions have been reported in patients diagnosed with atypical CML, chronic myeloproliferative neoplasm (cMPN), acute myeloid leukemia (AML), or ALL (*5*–*18*). TEL-ABL fusion oncogenes produce two different TEL-ABL proteins (type A and type B), depending on the length of the TEL sequence involved in the translocation.

Structural analyses of c-Abl have helped explain why the fusion proteins have unregulated kinase activity. The N-terminal segment of c-Abl includes a Src-homology 3 (SH3) domain, a Src-homology 2 (SH2) domain, and a tyrosine kinase domain (Figure 1A and 1B). The SH3-SH2-kinase unit (“Src-module”) is the core signaling and catalytic unit of c-Abl. The Src-module of c-Abl is followed by a 615 residue segment that is dispensible for the regulated kinase activity of ABL (*19*). The Src-module of c-Abl is preceded by an N-terminal cap (N-Cap) segment that is myristoylated at the very N-terminus. c-Abl is autoinhibited by the binding of the myristoyl group to a deep hydrophobic pocket at the base of the kinase domain (*20*, *21*). Myristoyl binding induces a conformational change in the kinase domain that allows SH2 docking onto the distal surface of the C-lobe of the kinase. Specifically, the C-terminal helix of the kinase domain (*α*I) bends sharply upon myristoyl binding, resulting in hydrophobic contacts between the bent portion of the helix (*α*I’) and the myristoyl group. This, in turn, enables docking of the SH3 domain on the linker connecting the SH2 domain to the N-lobe of the kinase domain, and these interactions suppress kinase activity (Figure 1B) (*20*, *22*).

Both BCR-ABL and TEL-ABL proteins contain self-association domains that promote oligomerization, resulting in kinase activation by trans-autophosphorylation (*7*, *23*– *25*). In BCR-ABL, a coiled-coil domain mediates self-association, most likely resulting in dimers (*25*–*27*). A pointed (PNT) domain in TEL forms an open spiral oligomer by itself, and so likely drives the formation of dimers and higher-order oligomers of TEL-ABL (*7*, *28*, *29*). Since the ABL component of the fusion protein is located after the fusion partner, the very N-terminal segment of ABL, including the myristoylation site, is lost upon formation of the fusion protein, leading to disruption of the myristoyl-mediated kinase autoinhibition mechanism.

A breakthrough in the treatment of CML was the development of the BCR-ABL inhibitor imatinib (Gleevec), which was the first kinase inhibitor approved for clinical use (*30*). Imatinib and more recent ATP-competitive ABL inhibitors, such as dasatinib (Sprycel) (*31*, *32*), nilotinib (Tasigna, AMN107) (*33*), and ponatinib (Iclusig) (*34*), are remarkably effective, but suffer from the emergence of resistance mutations in patients. An important advance in this regard was the development of asciminib (Scemblix, ABL001), an allosteric inhibitor that binds to the myristoyl-binding pocket in the kinase domain of ABL (*35*, *36*). Asciminib acts as a surrogate for the myristoyl group, and it induces the same conformational change in the C-terminal helix of the kinase domain (*α*I) that is induced by binding of the myristoyl group (*36*). Thus, the binding of asciminib repairs the broken autoinhibitory switch in the kinase domain of the fusion protein, enabling the docking of the SH2 domain on the C-lobe of the kinase domain. These interactions suppress the catalytic activity of the kinase by stabilizing the same inactive conformation of the Src-module that is seen in myristoylated and auto-inhibited c-Abl (*36*). Because asciminib binds at a site that is distant from the active site, it can be used in combination with ATP-competitive inhibitors such as imatinib, nilotinib, and ponatinib, with improved therapeutic effects (*37*–*41*). This combination enables suppression of otherwise untreatable compound mutants of BCR-ABL at clinically achievable concentrations and also suppresses the emergence of resistance mutations in cell models (*37*–*41*).

Imatinib and other ATP-competitive inhibitors are similarly effective against both p210 BCR-ABL and TEL-ABL, which is expected since the ABL components (corresponding to residues 28 to 1130 in ABL1a) are identical in sequence in the two fusion proteins (Figure 1A). Unexpectedly, GNF-2, a BCR-ABL inhibitor that was a precursor to the development of asciminib, is ineffective against TEL-ABL (*42*). GNF-2 binds to the myristoyl pocket of the ABL kinase domain similarly to asciminib (*43*, *44*).

We were intrigued by the unexpected resistance of TEL-ABL to GNF-2, and so we evaluated the ability of asciminib to inhibit BCR-ABL and TEL-ABL in cells. To do this, we monitored the effects of these compounds on the growth of Ba/F3 or 32D cells transformed by p210 BCR-ABL (referred to hereafter as BCR-ABL) or TEL-ABL type A (referred to hereafter as TEL-ABL). We found that asciminib is a potent inhibitor of cell growth induced by BCR-ABL, with an IC_50_ value in the nanomolar range, consistent with previous results (*35*, *36*, *45*, *46*). Surprisingly, we found that asciminib was >2000-fold less potent at inhibiting cell growth driven by TEL-ABL, with an IC_50_ value in the micromolar range. The behavior of asciminib is in marked contrast to that of ATP-competitive inhibitors, such as ponatinib and imatinib, which are nanomolar inhibitors of cell proliferation driven by either BCR-ABL or TEL-ABL.

We wondered whether differences in the oligomerization properties of the TEL and BCR components of the fusion proteins could be responsible for differences in the sensitivity to asciminib, through differences in the levels of autophosphorylation. To quantify the oligomerization and phosphorylation of BCR-ABL and TEL-ABL, we used a single-molecule imaging assay in which ABL fusion proteins are tagged with an N-terminal fluorescent protein, expressed and biotinylated in mammalian cells, and then captured on streptavidin-coated glass slides. By quantitative analysis of fluorescence intensity from individual molecular complexes, we found that BCR-ABL is mainly dimeric, but that TEL-ABL forms higher-order oligomers, such as tetramers, hexamers, and higher-order species. We monitored tyrosine phosphorylation of both TEL-ABL and BCR-ABL fusion proteins using a non-specific anti-phosphotyrosine antibody, 4G10, as well as site-specific antibodies that recognize phosphorylation on the activation loop (Tyr 412 in ABL1b numbering) of the kinase domain, or phosphorylation of two important regulatory tyrosine residues in the SH3 domain (Tyr 89 in ABL1b numbering) (*47*) or in the SH2-kinase linker (Tyr 245 in ABL1b numbering) (*48*). We found that Tyr 89 phosphorylation levels, in particular, were markedly higher in TEL-ABL compared to BCR-ABL.

The single-molecule assay was implemented in flow cells, which allowed BCR-ABL and TEL-ABL to be captured from cell lysates, immobilized in the flow cell, and then treated with different reaction conditions, such as the addition of ATP-Mg^2+^, or incubation with YopH, a tyrosine phosphatase, as well as the ABL inhibitors, asciminib and ponatinib. Using this assay, we found that when TEL-ABL is phosphorylated, it is insensitive to asciminib. However, when TEL-ABL is dephosphorylated by a phosphatase, asciminib sensitivity is restored. These results lead us to conclude that the formation of higher-order oligomers promotes enhanced tyrosine phosphorylation of TEL-ABL and causes resistance to asciminib.

## RESULTS and DISCUSSION

### TEL-ABL is resistant to the allosteric inhibitor asciminib

Ba/F3 cells are murine pro-B cells that rely on interleukin-3 (IL-3) for survival and proliferation. Upon transduction of oncogenic proteins such as BCR-ABL and TEL-ABL, Ba/F3 cells can grow in the absence of IL-3 (*49*). Addition of inhibitors of the oncogenic proteins attenuates cell growth. We compared the effects of the allosteric ABL inhibitor asciminib (*35*) with that of a recently developed ATP-competitive inhibitor, ponatinib (*34*). Ba/F3 cells were retrovirally transduced with BCR-ABL or TEL-ABL constructs. Following transduction, IL-3 was withdrawn from the growth medium. These cells were then treated with varying concentrations of either asciminib or ponatinib and cell growth was measured by flow cytometry. Cells expressing BCR-ABL were sensitive to asciminib (IC_50_: *∼*2 nM), as shown previously (*35*). In contrast, asciminib was >2000-fold less potent at inhibiting the growth of Ba/F3 cells expressing TEL-ABL (IC_50_: *∼*13 μM) (Figure 2A, middle panel). Ponatinib, on the other hand, was effective at nanomolar concentrations against both ABL fusion proteins (BCR-ABL IC_50_: *∼*4 nM, TEL-ABL IC_50_: *∼*4 nM) (Figure 2B, middle panel).

**Fig. 2.**
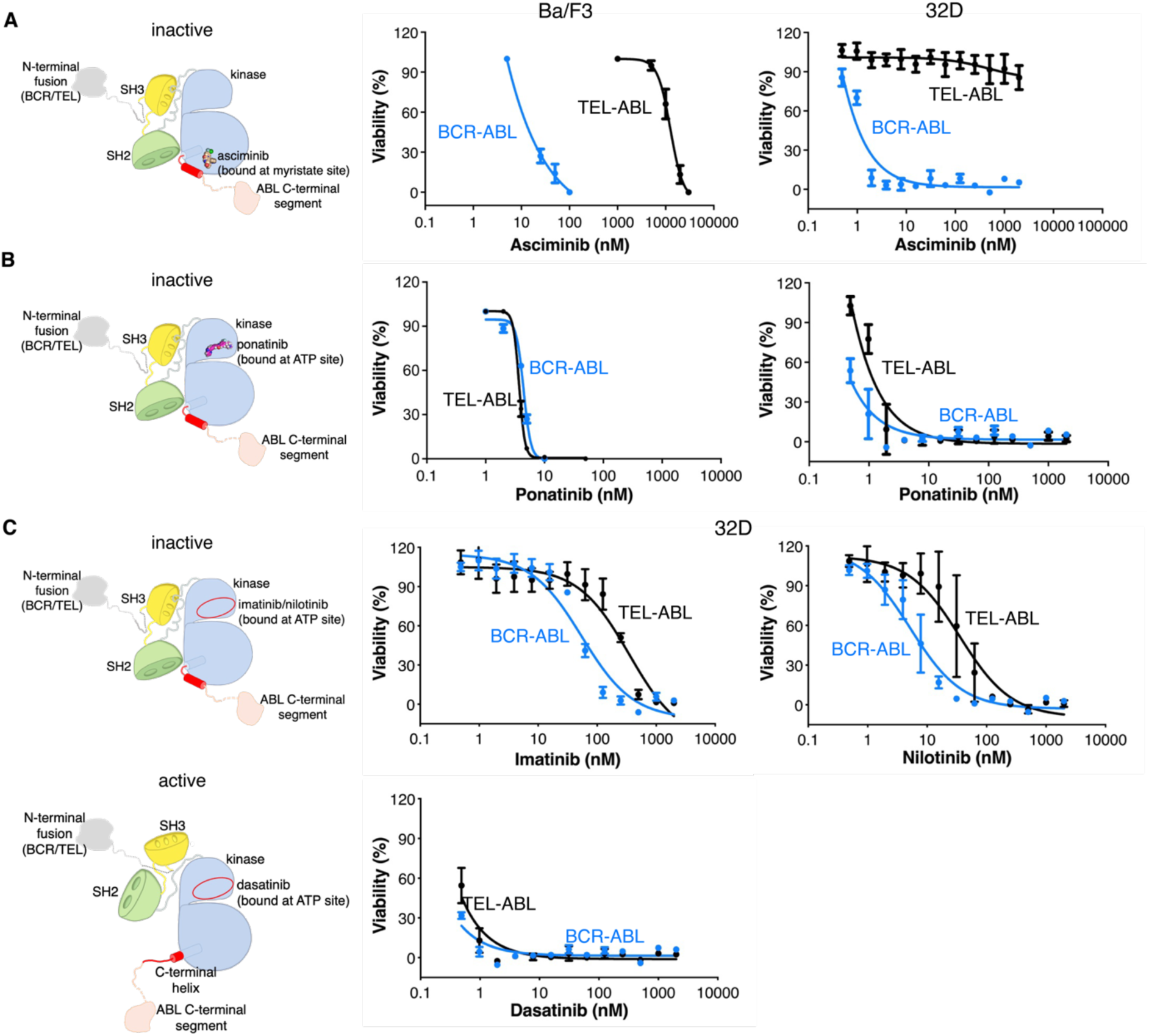
TEL-ABL is resistant to the allosteric inhibitor asciminib. **(A**) Dose-response curves for asciminib in Ba/F3 and 32D cell lines. Schematic representations of fusion proteins bound to the inhibitors are shown on the left. Data points represent the mean percent of untreated ± SEM. **(B)** Dose-response curves for ponatinib in Ba/F3 and 32D cell lines. Data points represent the mean percent of untreated ± SEM. (**C)** Dose-response curves for imatinib, nilotinib, and dasatinib in 32D cell line. Data points represent the mean percent of untreated ± SEM (N=3, number of experiments).

The 32D murine myeloid cell line is also dependent on IL-3 for survival and growth (*50*, *51*). Introduction of BCR-ABL or TEL-ABL transforms 32D cells, and they become independent of IL-3 (*52*). 32D cells that were deprived of IL-3 were treated with varying concentrations of asciminib and the ATP-competitive inhibitors imatinib, nilotinib, dasatinib, and ponatinib. As for Ba/F3 cells, 32D cells expressing BCR-ABL were sensitive to asciminib (IC_50_: *∼*1 nM). Cells expressing TEL-ABL were resistant to asciminib (IC_50_ > 2 μM; Figure 2A, right panel). In contrast, all ATP-competitive inhibitors tested were effective at inhibiting the growth of 32D cells expressing either BCR-ABL or TEL-ABL (Figure 2B-D). All subsequent analyses are focused on the comparison of asciminib with a single ATP-competitive inhibitor, ponatinib.

### The SH2 and SH3 domains do not appear to exert inhibitory control in TEL-ABL

A key aspect of the inhibition of BCR-ABL by asciminib is the stabilization of the bent conformation of helix *α*I by asciminib (*36*). The bent conformation arises from a localized kink around Met 496 of helix *α*I, such that the last 12 residues (Ser 501-Lys 512 in ABL1a numbering), denoted *α*I’, are oriented orthogonally to the main body of the helix. Helix αI′ is critical for asciminib binding, and it gates SH2 docking.

To see whether asciminib alters the C-terminal helix conformation in the same way in TEL-ABL as in BCR-ABL, we sought to study the effects of deleting helix *α*I’. To simplify the deletion constructs, we first established that deleting the C-terminal segment of the ABL component (corresponding to residues 582 to 1130 in ABL1a) does not affect the activity and drug sensitivity of either BCR-ABL or TEL-ABL. We refer to the ABL component lacking the C-terminal segment as ABL(core). Both BCR-ABL(core) and TEL-ABL(core) fusion proteins conferred IL-3–independence in Ba/F3 cells (Figure S1). These Ba/F3 lines were then treated with ponatinib or asciminib and cell viability was assessed. The drug sensitivity profiles of ABL(core) fusion proteins were the same as that of the intact fusion proteins. Asciminib inhibited Ba/F3 cells expressing BCR-ABL(core) potently (IC_50_: *∼*6 nM), but was >2000 fold less effective at inhibiting cells expressing TEL-ABL(core) (IC_50_: *∼*11 μM) (Figure 3A). Ponatinib inhibited the growth of cells expressing either BCR-ABL(core) or TEL-ABL(core) potently (IC_50_ BCR-ABL(core): *∼*6 nM, IC_50_ TEL-ABL(core): *∼*4 nM) (Figure 3B).

**Fig. 3.**
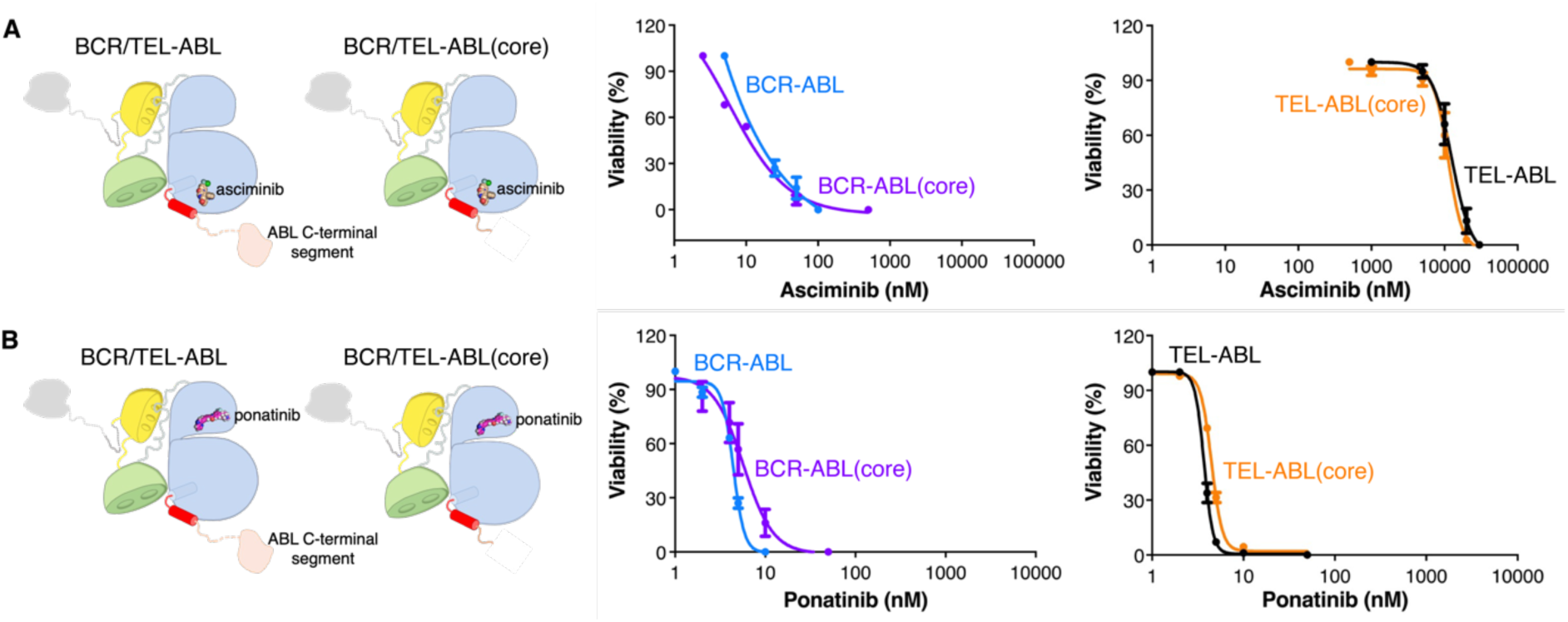
Deletion of the C-terminal segment of the ABL component does not affect the drug sensitivity of either BCR-ABL or TEL-ABL. **(A)** Dose-response curves for asciminib in Ba/F3 cell line. BCR-ABL/TEL-ABL: full-length BCR-ABL and TEL-ABL, including the entire ABL component, without truncation; BCR-ABL(core)/TEL-ABL(core): fusion proteins with ABL component lacking the C-terminal segment. Schematic representations of fusion proteins bound to the inhibitors are shown on the left. Data points represent the mean percent of untreated ± SEM. **(B)** Dose-response curves for ponatinib in Ba/F3 cell line. Data points represent the mean percent of untreated ± SEM (N=3, number of experiments).

We deleted helix αI’ in BCR-ABL(core) and TEL-ABL(core) (we refer to the ABL component of these deletion constructs as ABL(core*Δ*I’)) (Figure 4A), and performed proliferation assays using Ba/F3 cells. Ponatinib effectively inhibited both BCR-ABL (core*Δ*I’) and TEL-ABL(core*Δ*I’) fusion proteins, with IC_50_ values ranging between *∼*1 and *∼*6 nM (Figure 4C). Cells expressing BCR-ABL(core*Δ*I’) were not sensitive to asciminib (IC_50_: *∼*9 μM) (Figure 4B). This result, interpreted in the light of the structure of ABL1a (*21*, *22*), underscores the importance of helix αI’ in gating SH2-SH3 docking in asciminib-induced inhibition. In contrast, deletion of αI’ in TEL-ABL(core) did not alter the already high IC_50_ value for asciminib inhibition (Figure 4B). This result suggests that the gating provided by helix αI’ is inoperative in TEL-ABL, implying that the SH2 and SH3 domains no longer exert inhibitory control.

**Fig. 4.**
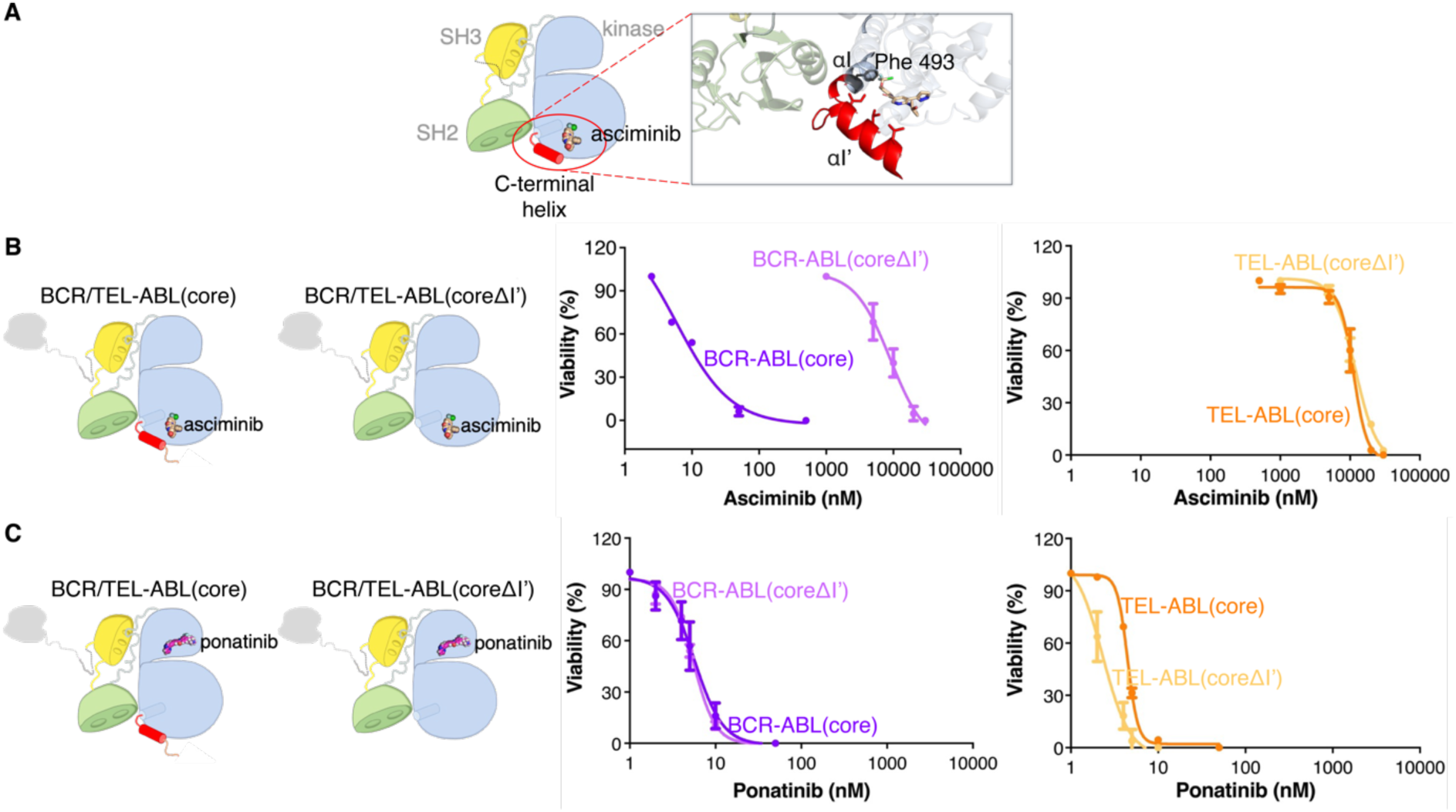
The SH2 and SH3 domains do not appear to exert inhibitory control in TEL-ABL. **(A)** Schematic representation of Abl “Src-module” bound to asciminib. In the zoom window, a cartoon representation of the crystal structure of asciminib-bound Abl is illustrated (PDB ID: 5MO4). Asciminib is shown as orange sticks. The distal lobe of the C-terminal helix is shown in red. Residues located on helix *α*I’ and important for asciminib binding are shown as red sticks. Phe 493 (shown as a gray stick) is the last residue in ABL(core*Δ*I’) fusion proteins. **(B)** Dose-response curves for asciminib in Ba/F3 cell line. BCR-ABL(core)/TEL-ABL(core): fusion proteins with ABL component lacking the C-terminal segment; BCR-ABL(core*Δ*I’)/TEL-ABL(core*Δ*I’): fusion proteins with ABL component lacking the helix *α*I’ (shown in red) in addition to the C-terminal segment. Schematic representations of fusion proteins bound to the inhibitors are shown on the left. Data points represent the mean percent of untreated ± SEM. **(C)** Dose-response curves for ponatinib in Ba/F3 cell line. Data points represent the mean percent of untreated ± SEM. (N=3, number of experiments).

### TEL-ABL forms higher-order oligomers than BCR-ABL

We designed a flow-cell assay that enables quantification of oligomerization of ABL fusion proteins using single-molecule Total Internal Reflection Fluorescence (TIRF) microscopy (*53*, *54*). In this and all subsequent experiments, we used full-length BCR-ABL and TEL-ABL, including the entire ABL component, without truncation. We fused a fluorescent protein (mNeon Green, mNG) (*55*) to the N-terminus of TEL-ABL or BCR-ABL. The fluorescent fusion proteins also contained a biotinylation sequence (Avitag) (*56*) at the N-terminus and were expressed in HEK 293T cells.

The co-expression of the *E. coli* biotin ligase, BirA, in HEK 293T cells results in the biotinylation of TEL-ABL and BCR-ABL. Following lysis, the cell lysate was diluted in Dulbecco’s Phosphate-Buffered Saline (DPBS) and added to a streptavidin-coated well in the flow chamber for 1 min, before washing it out with DPBS. During this incubation period, the biotinylated mNG-ABL fusion proteins were captured on the surface of the functionalized glass substrates through streptavidin-biotin interactions (see Materials and Methods) (Figure 5A).

**Fig. 5.**
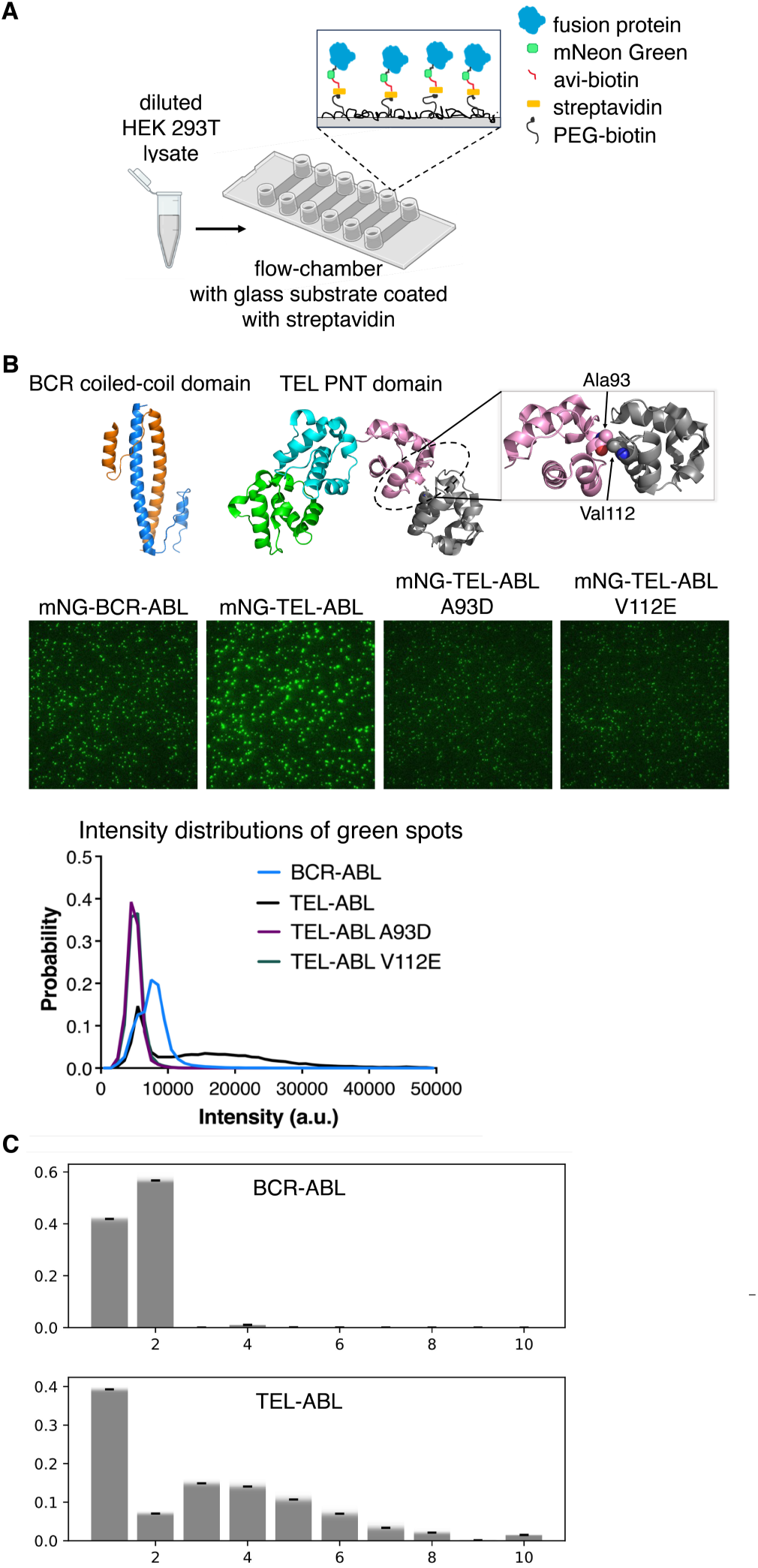
TEL-ABL forms higher-order oligomers than BCR-ABL. **(A)** Schematic diagram of mammalian expression-based single-molecule Total Internal Reflection Fluorescence (TIRF) assay. Biotinylated mNG-ABL fusion proteins overexpressed in HEK 293T cells were pulled down directly from diluted cell lysate, allowing visualization at a single-molecule resolution. The immobilization onto glass substrates functionalized with streptavidin relies on the interaction between biotinylated fusion protein and streptavidin. **(B)** Crystal structures of BCR oligomerization domain, coiled-coil domain (PDB ID: 1K1F), and TEL PNT domain (PDB ID: 7JU2) are shown on the top. The surface representation of the TEL PNT domain tetramer is shown. In the zoom window, residues located at the PNT oligomerization interface, A93 and V112E, are shown in spheres. Representative single-molecule TIRF images showing wild-type and monomeric mNG-ABL fusion proteins (green dots) are shown in the middle. Distribution of intensity for mNG-BCR-ABL, mNG-TEL-ABL, and monomeric mNG-TEL-ABL mutants (mNG-TEL-ABL A93D and mNG-TEL-ABL-V112E) (at 488 nm) are plotted on the bottom left. N=3 (number of experiments, only the mean intensity distribution is plotted) **(C)** Approximate oligomeric state distributions for BCR-ABL and TEL-ABL, extracted from the intensity distributions shown in (B) and (Fig S3), are plotted as bar graphs. Deconvolution of the measured instensity distribution data was performed using a lognormal approximation for single fluorophore intensity distributions and a moment-matched lognormal approximation for their summed intensities as well as a background correction.

The single-molecule fluorescence assay reveals the differing oligomeric states of BCR-ABL and TEL-ABL. HEK 293T cells expressing biotinylated mNG-BCR-ABL or mNG-TEL-ABL were harvested and lysed. The lysate was flowed over the streptavidin-coated glass surface and the captured complexes were imaged using TIRF. The mNG-fusion proteins were detected with 488 nm laser excitation (green channel). The histograms of fluorescence intensities revealed very different profiles for BCR-ABL and TEL-ABL. For BCR-ABL, there is a peak in the intensity distribution, centered at *∼*8*×*10^3^ intensity units (arbitrary intensity scale) and there is a resolved shoulder to this peak, at *∼*5*×*10^3^ intensity units. For TEL-ABL, there is a peak at *∼*5*×*10^3^ intensity units, corresponding to the shoulder in the BCR-ABL distribution, and a broad distribution of higher intensity values, ranging up to *∼* 30×10^3^ intensity units (Figure 5B).

The oligomerization of the isolated PNT domain in TEL-ABL can be disrupted by the introduction of point mutations located at the oligomerization interface (A93D and V112E) (*29*) (Figure 5B). We generated variants of TEL-ABL fusion proteins with these mutations in the PNT domain. HEK 293T cells expressing these TEL-ABL mutants were lysed and the fusion proteins were captured on a streptavidin-coated glass surface and imaged using TIRF. The distribution of fluorescence intensities revealed a single peak at low fluorescence intensity (*∼*5*×*10^3^ intensity units) for each of the TEL-ABL mutants. This confirms that the crystallographically observed interfaces between PNT domains are relevant for the oligomerization of TEL-ABL, and also suggests that the peak or shoulder at *∼*5*×*10^3^ intensity units for the BCR-ABL and TEL-ABL distributions correspond to the monomeric fluorescent protein (Figure 5B). Using step photobleaching analyses, we verified that these peaks at lower fluorescence intensity do indeed correspond to monomers (Figure S2A and Figure 5B). This analysis demonstrates that TEL-ABL forms higher-order oligomers than BCR-ABL. We next deconvolved the measured intensity distributions to obtain approximate distributions of oligomeric species for the two fusion proteins. For this analysis, we used a lognormal approximation for the observed fluorescence intensity distribution from a single fluorophore, and similarly use a lognormal approximation for the summed intensity of multiple fluorophores, in line with previous studies (61) and the appearance of our data (Figure S3) (see Materials and Methods).

Analysis of fluorescence intensity profiles demonstrates that BCR-ABL is mostly monomeric (*∼*40%) or dimeric (*∼*55%), with a small (*∼*5%) tetramer population. In contrast, TEL-ABL had a substantial multimer population (Figure 5C, Figure S2B, Figure S3). We observed trimers, tetramers, hexamers, and higher-order species in the fluorescence intensity profile of TEL-ABL (Figure 5C and Figure S2C). This is consistent with the crystal structure of the PNT domain, which shows that it forms a helical open-ended oligomer (*28*, *29*).

### TEL-ABL phosphorylation is not affected by asciminib in HEK 293T cells

We used the flow-cell assay to measure the autophosphorylation of ABL fusion proteins under different reaction conditions using single-molecule TIRF microscopy (*53*, *54*). mNG-BCR-ABL and mNG-TEL-ABL fusion proteins were expressed in HEK 293T cells, which were then treated with dimethyl sulfoxide (DMSO) solutions of inhibitors (asciminib or ponatinib) or a reference solution containing DMSO alone. Following inhibitor treatment, cells were harvested and lysed.

The mNG-fusion proteins were detected with 488 nm laser excitation (green channel). Phosphorylation levels were monitored using either a non-specific pan-phosphotyrosine antibody or a site-specific phospho-Tyr89 antibody. The antibodies were labeled with Alexa-568 and detected with 561 nm laser excitation (red channel). Representative two-color TIRF images from these experiments are shown in Figure 6A.

**Fig. 6.**
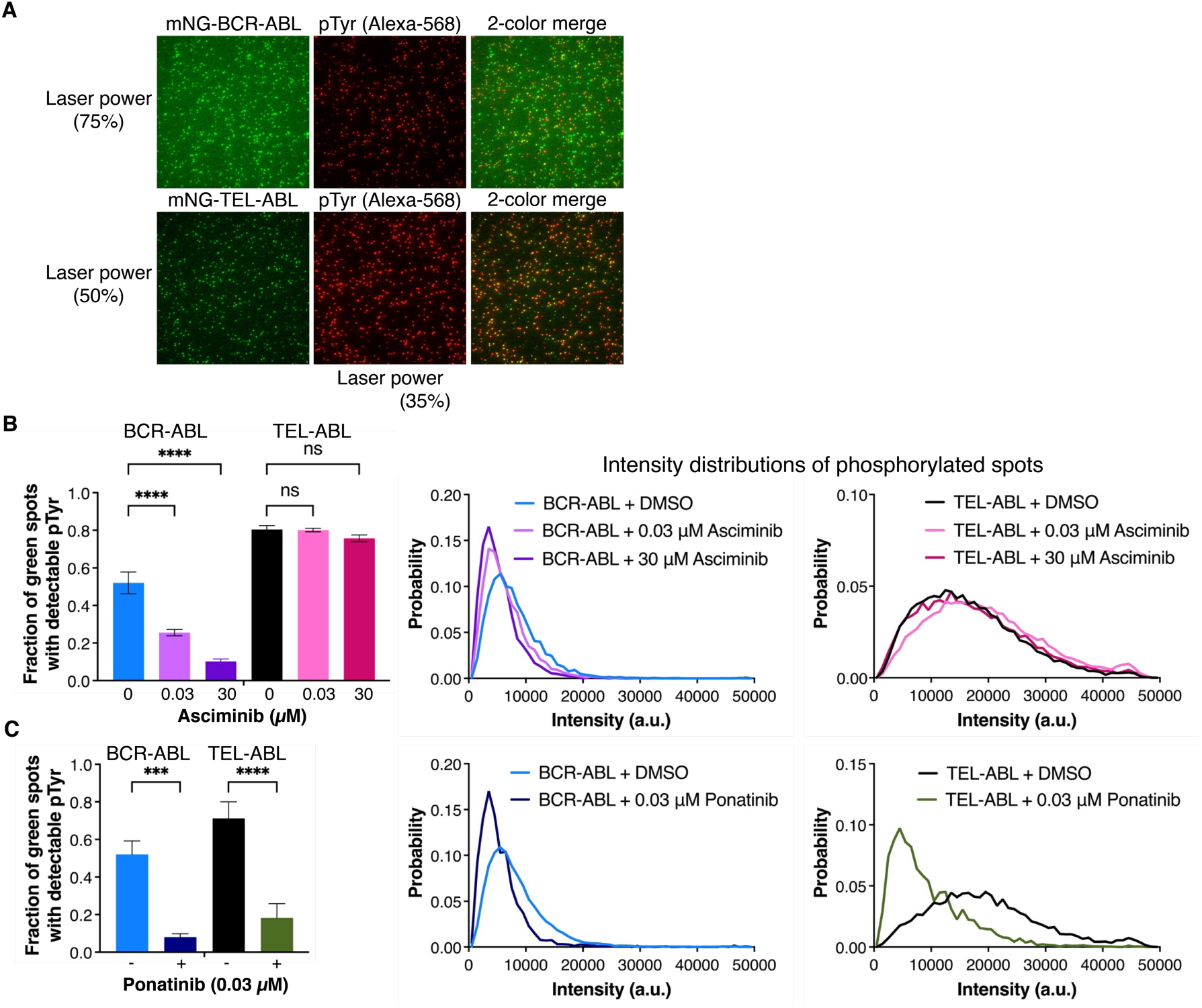
TEL-ABL phosphorylation is not affected by asciminib in HEK 293T cells. **(A)** Representative single-molecule TIRF images showing mNG-ABL fusion proteins (green dots), total tyrosine phosphorylation (red dots), and a 2-color merge of these images reporting on the fraction of phosphorylated ABL fusion spots are shown on the right. **(B)** The fraction of mNG-ABL fusion proteins that show detectable tyrosine phosphorylation is plotted for different asciminib concentrations (0.03 μM and 30 μM). HEK 293T cells treated with DMSO or inhibitor were harvested, lysed, and pulled down onto a coated glass surface. The autophosphorylation status of mNG-ABL fusion proteins was measured using pan-pTyr antibody and Alexa-568 labeled secondary antibody. The distribution of intensity for pTyr (561 nm), at different asciminib concentrations, for mNG-ABL fusion proteins with detectable phosphorylation are plotted on the right (see Materials and Methods for details of normalization). **(C)** The fraction of mNG-ABL fusion proteins that show detectable tyrosine phosphorylation is plotted for 0.03 μM ponatinib. The distribution of intensity for pTyr (561 nm), at 0.03 μM ponatinib, for mNG-ABL fusion proteins with detectable phosphorylation are plotted on the right (see Materials and Methods for details of normalization). (One-way ANOVA, N=3, number of experiments, only the mean intensity distribution is plotted) (**** p*≤*0.0001, *** p*≤*0.001, ** p*≤*0.01, *p*≤*0.05, ns p>0.05).

We quantified the extent of phosphorylation of the fusion proteins in two ways. First, we measured the fraction of green fluorescent spots, corresponding to mNG-labeled ABL fusion protein complexes, that are colocalized with red spots (Alexa-568 labeled antibodies reporting on phosphorylation). This provides a measure of fusion protein phosphorylation. After asciminib treatment, we observed a decrease in the fraction of BCR-ABL spots that are phosphorylated. In contrast, for TEL-ABL, asciminib treatment results in no detectable change in the fraction of spots that are phosphorylated, indicating that asciminib does not inhibit the kinase activity of TEL-ABL effectively in cells (Figure 6B, left panel).

We also quantified the mean level of phosphorylation per individual spot, by integrating the intensity distributions for the red channel (corresponding to phosphotyrosine levels) and normalizing by the total number of green spots detected. Individual ABL fusion complexes with a larger number of phosphorylated tyrosine residues would bind more phospho-specific antibodies, which leads to enhancement in the signal intensity (a right shift in the intensity distribution of the red channel). A limitation of this assay is that we cannot rely on the integrated intensity values to determine the absolute number of phosphate groups within each complex, because it is not possible to know whether every phosphate group has an antibody bound to it. Steric interference between antibodies, or interference with the glass support, can affect antibody binding. Due to this limitation, we use the values of the integrated antibody fluorescence intensities to provide a relative, rather than an absolute, measure of the extent of phosphorylation.

A slight reduction in the extent of phosphorylation was observed in BCR-ABL treated with asciminib, but phosphorylation levels remained unchanged in TEL-ABL treated with asciminib (Figure 6B, right panel). Ponatinib effectively inhibited both BCR-ABL and TEL-ABL, as evidenced by a decrease in the fraction of phosphorylated spots and the extent of phosphorylation per spot (left shift upon ponatinib treatment) (Figure 6C).

### TEL-ABL spots have more detectable phosphorylation on Tyr 89 compared to BCR-ABL

We asked whether differences in the oligomeric states of TEL-ABL and BCR-ABL affect the levels of phosphorylation of the activation loop (Tyr 412) or the two other sites of regulatory tyrosine (Tyr 89 and Tyr 245) phosphorylation (Figure 7A, left panel). Although all three sites are expected to be phosphorylated (*47*, *48*, *57*), appreciable levels of phosphorylation in the flow cell assay were only detected for Tyr 89, so we focused on this residue for the analysis. Limited detection of Tyr 245 and Tyr 412 may reflect interactions of the captured proteins with the glass, or the fact that the proteins in this assay are folded, as opposed to the unfolded nature of the samples in conventional western blot assays.

**Fig. 7.**
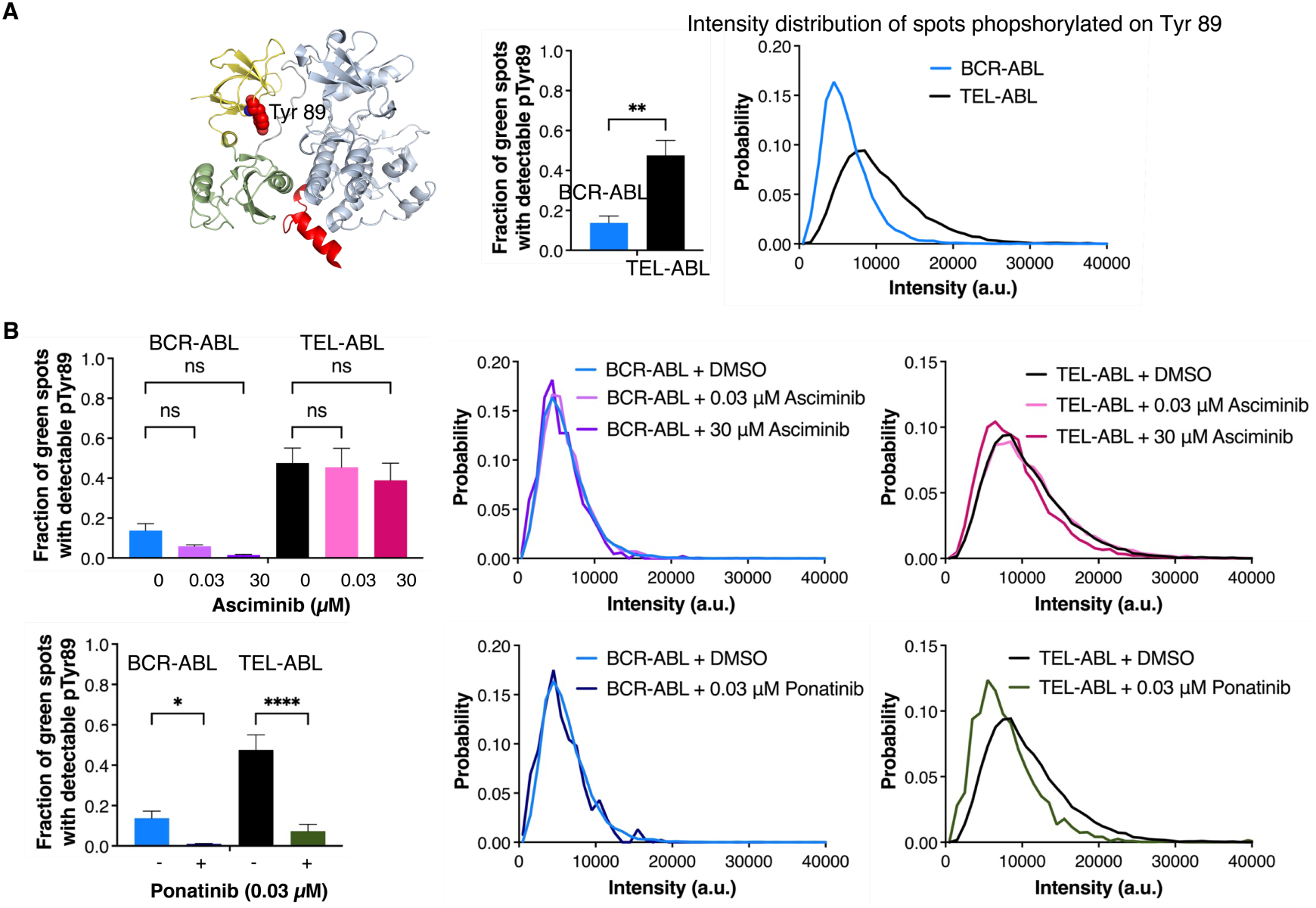
TEL-ABL spots show more detectable phosphorylation on Tyr 89 compared to BCR-ABL. **(A)** Crystal structure of autoinhibited ABL “ Src-module” showing Tyr 89, located on the SH3 domain, in red spheres. The fraction of mNG-ABL fusion proteins that show detectable Tyr 89 phosphorylation is plotted. The distribution of intensity for phospho-Tyr 89 (at 561 nm), for mNG-BCR-ABL and mNG-TEL-ABL fusion proteins with detectable phosphorylation are plotted on the right. **(B)** The fraction of mNG-ABL fusion proteins that show detectable Tyr 89 phosphorylation is plotted for different asciminib (0.03 μM and 30 μM) and ponatinib (0.03 μM) concentrations. (One-way ANOVA). The distribution of intensity for pTyr 89 (561 nm), at different asciminib and ponatinib concentrations, for mNG-ABL fusion proteins with detectable phosphorylation are plotted on the right. N=3 (number of experiments, only the mean intensity distribution is plotted) (**** p*≤*0.0001, *** p*≤*0.001, ** p*≤*0.01, *p*≤*0.05, ns p>0.05).

We found that the fraction of TEL-ABL spots phosphorylated on Tyr 89 was significantly greater than for BCR-ABL. Whereas *∼*50% of the fluorescent spots corresponding to TEL-ABL showed detectable phosphorylation at Tyr 89, only *∼*10% of the BCR-ABL spots did so (Figure 7A, middle panel). TEL-ABL also had a higher degree of Tyr 89 phosphorylation per spot as evident by the peak at higher fluorescence intensity (Figure 7A, right panel).

Given that TEL-ABL forms larger assemblies than BCR-ABL, it is not surprising that spots corresponding to TEL-ABL are more likely to contain phosphorylated residues. Our data do not have the precision to determine whether per-molecule phosphorylation levels are higher for TEL-ABL than for BCR-ABL. Instead, we simply conclude from the data that the larger TEL-ABL assemblies are more likely to contain phosphorylated fusion proteins than the smaller BCR-ABL assemblies. Since phosphorylation on Tyr 89 results in kinase activation, we reason that assemblies containing some phosphorylated species can promote transphosphorylation and activation of other proteins in the assembly, leading to maintenance of higher levels of kinase activity, particularly in the face of dephosphorylation by phosphatases.

We examined the levels of Tyr 89 phosphorylation in BCR-ABL and TEL-ABL when HEK 293T cells expressing these fusion proteins are treated by asciminib or ponatinib. We found that the fraction of TEL-ABL spots that are phosphorylated and the extent of phosphorylation per spot were not affected by asciminib treatment of the cells. In contrast, for cells expressing TEL-ABL and treated with ponatinib, there was little detectable Tyr 89 phosphorylation. The fraction of BCR-ABL spots with detectable phosphorylation on Tyr 89 was decreased by both ponatinib and asciminib treatment (Figure 7B, left panels). For those green spots that are phosphorylated, we did not observe any substantial decrease in the fluorescence intensity distribution of the phospho-antibody (red channel) in BCR-ABL upon treatment with either inhibitor (Figure 7B, right panels). Very dim spots (e.g. spots with single phosphotyrosine residues) may not be detected by the assay. This reduces the dynamic range of the assay for BCR-ABL, for which only one or two phospho-Tyr 89 residues can be present.

We also measured the phosphorylation levels of monomeric TEL-ABL mutants in HEK 293T cells. Mutational disruption of the predicted PNT domain interface in TEL-ABL led to a marked decrease in the fraction of phosphorylated TEL-ABL spots and the extent of phosphorylation per spot (Figure S4). We next tested the ability of these monomeric TEL-ABL mutants to confer IL-3 independence in Ba/F3 cells. As expected, cells transduced with these mutants could not grow in the absence of IL-3 (data not shown).

### TEL-ABL retains an intrinsic capacity to be inhibited by asciminib

Using the single-molecule assay, we asked whether dephosphorylation of TEL-ABL can restore asciminib sensitivity. TEL-ABL and BCR-ABL fusion proteins captured on glass from HEK 293T cells were incubated with the tyrosine phosphatase YopH (*58*). The flow cells were then washed with the gel filtration buffer to remove the phosphatase, followed by the addition of inhibitors (ponatinib or asciminib) (See Materials and Methods). After a defined period of inhibitor treatment, the flow cells were washed with the gel filtration buffer to remove the inhibitor. Kinase reactions were then initiated by flowing in ATP-Mg^2+^. The autophosphorylation status of both fusion proteins after the reaction was determined using the pan-phosphotyrosine antibody (4G10) and the specific phospho-Tyr 89 antibody.

YopH phosphatase was effective at dephosphorylating both TEL-ABL and BCR-ABL captured from cells, as demonstrated by a reduction in the fluorescence intensity corresponding to the phosphotyrosine antibodies (Figure 8A). Tyr 89 phosphorylation was almost completely removed in both ABL fusion proteins, as shown by the substantial reduction in the fraction of phosphorylated spots (Figure S4A). For both TEL-ABL and BCR-ABL, incubation with ATP-Mg^2+^ restored Tyr 89 phosphorylation, as demonstrated by the shift in the peak maximum towards higher values and the increase in the fraction of spots that are phosphorylated on Tyr 89 (Figure 8A and Figure S5A). BCR-ABL and TEL-ABL pre-treated with either ponatinib or asciminib could not restore Tyr 89 phosphorylation following ATP-Mg^2+^ incubation. The distributions of antibody fluorescence intensities for these samples resemble that of YopH-treated samples (Figure 8A). This result shows that both BCR-ABL and TEL-ABL are sensitive to both ponatinib and asciminib when the proteins are dephosphorylated prior to inhibitor addition.

**Fig. 8.**
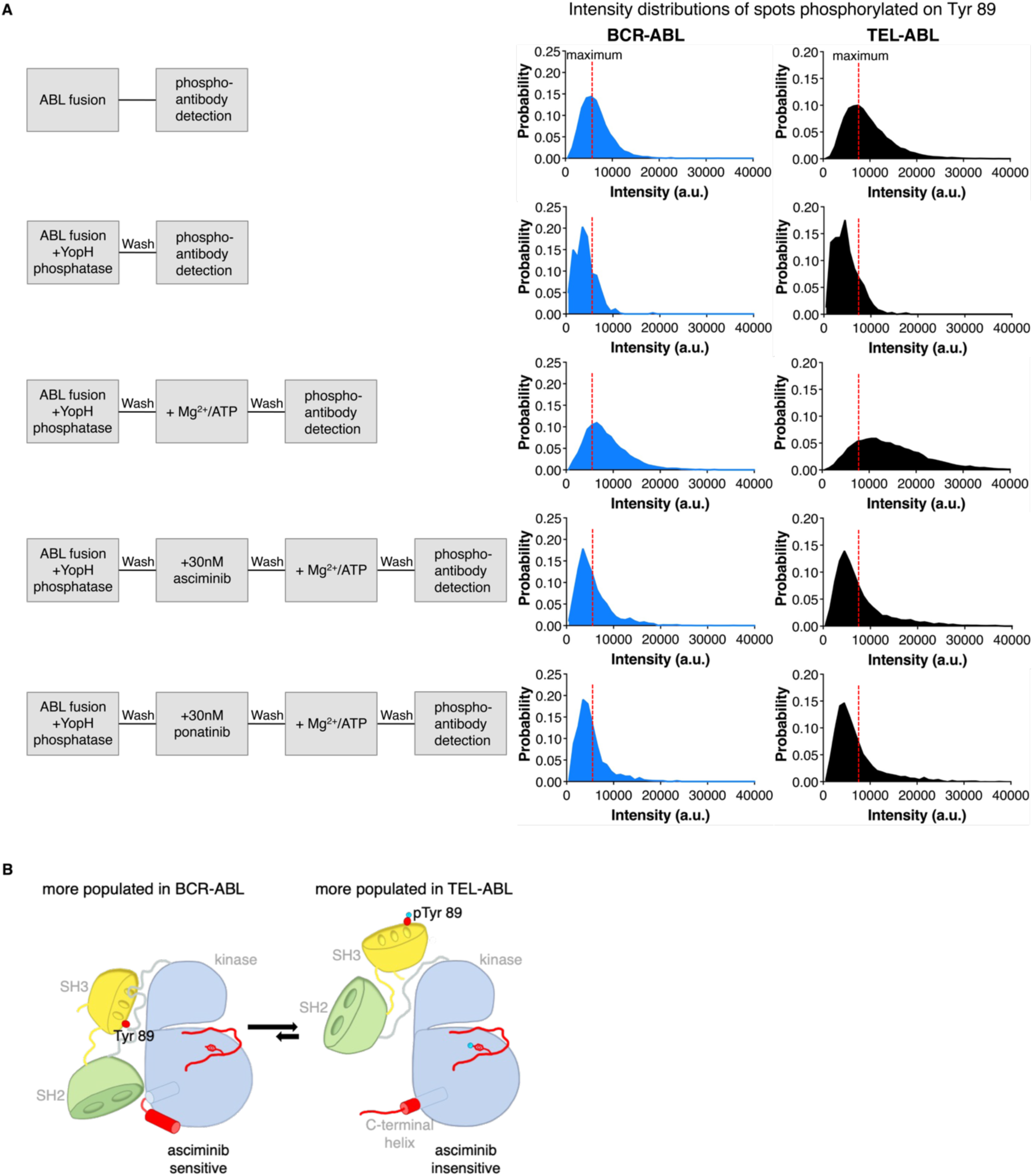
TEL-ABL retains an intrinsic capacity to be inhibited by asciminib. **(A**) A schematic diagram showing the experimental setup is shown on the left in the lower panel. The distribution of intensity for pTyr 89 (561 nm), at different reaction conditions, for mNG-ABL fusion proteins with detectable phosphorylation are plotted on the right. N=4 (number of experiments, only the mean intensity distribution is plotted). **(B)** The conformational equilibrium of BCR-ABL and TEL-ABL asciminib-sensitive (assembled, autoinhibited) and asciminib-insensitive (disassembled) states is shown. BCR-ABL and TEL-ABL are in a conformational equilibrium between the assembled and disassembled states, which is predominantly on the side of the assembled state for BCR-ABL and the disassembled state for TEL-ABL. Tyr 89 phosphorylation in the SH3 domain of ABL shifts this equilibrium to the disassembled, asciminib-insensitive state in TEL-ABL.

Qualitatively similar results were obtained when phosphorylation was monitored by the pan-phosphotyrosine antibody, 4G10. In this case, the fraction of spots that are phosphorylated was reduced only by 40% in YopH-treated samples (Figure S5B). We note that the co-localization assay does not distinguish between spots that have different levels of phosphorylation and the detection of even a single phosphotyrosine residue in the spot is treated the same as the detection of a higher level of phosphorylation. The appearance of a tail at higher fluorescence intensity values for the pan-phosphotyrosine antibody in the BCR-ABL distribution, and a second peak at a higher intensity in the TEL-ABL distribution suggest that both fusion proteins are active (Figure S5C). Evidence for kinase activity was absent in the samples treated with ponatinib or asciminib, after phosphatase treatment (Figure S4B). These data also support the conclusion that BCR-ABL and TEL-ABL are both intrinsically sensitive to ponatinib and asciminib when dephosphorylated.

## CONCLUDING REMARKS

Asciminib is the first allosteric inhibitor of BCR-ABL approved for use in the clinic (*35*, *59*), and it is effective against many BCR-ABL variants with mutations that confer resistance to ATP-competitive inhibitors (*45*, *46*). The special utility of asciminib arises from the fact that it binds to the kinase domain at a site that is far from the catalytic center. It is therefore not affected directly most of the time by mutation-induced changes to the active site and can be used in combination with ATP-competitive inhibitors.

Asciminib inhibits the kinase activity of BCR-ABL by mimicking the action of an N-terminal myristoyl modification of ABL1b, a natural allosteric modulator of ABL that is lost in oncogenic fusion variants. In this work, we compared the ability of asciminib and various ATP-competitive inhibitors (ponatinib, imatinib, nilotinib, and dasatinib) to inhibit BCR-ABL and TEL-ABL. The ABL component of BCR-ABL and TEL-ABL is identical in sequence and, as established previously, the ATP-competitive inhibitors are effective against both fusion proteins in cell-growth assays. Unexpectedly, while asciminib is a potent inhibitor of BCR-ABL, we discovered that it is >2000-fold less effective against TEL-ABL. We traced the origin of this differential sensitivity to differences in the levels of phosphorylation of TEL-ABL and BCR-ABL, including at a critical regulatory site in the SH3 domain of ABL, Tyr 89.

Asciminib binds to the myristoyl binding site in the C-lobe of the ABL kinase domain. The binding site is completed by the bending of the C-terminal helix of the kinase domain (*α*I) over either asciminib or the myristoyl group (*22*, *36*). Extensive hydrophobic interactions lead to mutual stabilization of the bound inhibitor and the bent conformation of helix *α*I. The bending of the helix is critical for SH2 docking onto the distal surface of the C-lobe of the kinase, which enables the assembly of the autoinhibited conformation of the Src module of ABL, which in turn stabilizes an inactive conformation of the catalytic center. This establishes a coupled equilibrium, by which destabilization of the internal docking of the SH2-SH3 assembly weakens the binding of asciminib or the myristoyl group to the kinase domain. Our data suggest that phosphorylation of ABL, which is known to disturb the docking of the SH2 and SH3 domains on the kinase domain, leads to resistance to asciminib through this coupled conformational equilibrium (Figure 8B). We also demonstrate that TEL-ABL is intrinsically susceptible to inhibition by asciminib – the observed resistance to asciminib of TEL-ABL isolated from mammalian cells can be reversed by treatment with a phosphatase.

All protein kinases are regulated by phosphorylation, and changes in oligomerization are the most common mechanism by which autophosphorylation rates are altered in normal signaling mechanisms. BCR-ABL and TEL-ABL are both oligomeric proteins, and it has long been appreciated that the oligomerization induced by the fusion partner potentiates autophosphorylation and unregulated activation. Our data show that TEL-ABL forms larger oligomers than does BCR-ABL. These findings are consistent with studies indicating that BCR protein forms antiparallel coiled-coil dimers (*26*) while the PNT domain of TEL forms an open-ended polymeric structure (*28*, *29*).

In a recent study published while we were preparing this manuscript, a rare BCR-ABL variant that lacks the myristoylated N-cap segment as well as the first 23 residues of the SH3 domain of ABL was shown to be resistant to asciminib in Ba/F3 cells (*60*). The SH3 domain of the ABL component of this BCR-ABL variant is unlikely to be folded, and the Src module cannot assemble in this variant. In agreement with our conclusions, this study points to the critical role played by the SH2 and SH3 domains in the autoinhibition of ABL and therefore for asciminib action. In this regard, note that asciminib stabilizes the assembled inactive state of Abl, while some ATP-competitive inhibitors, such as imatinib, stabilize a disassembled inactive state in which SH3-SH2 domains are detached from the kinase domain (*61*–*64*). This could explain why we do not see a difference in the sensitivity of BCR-ABL and TEL-ABL to inhibitors such as imatinib and ponatinib.

A key concept underlying the interpretation of our data is that protein kinases constantly switch between active and inactive states, with the relative population of these states determined by the balance between kinase and phosphatase action. Asciminib acts by stabilizing an inactive conformation of ABL, and even though the fusion proteins are active kinases, phosphatase action enables these proteins to sample this inactive state. Our data show that ATP-competitive inhibitors imatinib, nilotinib, and ponatinib are similarly effective against BCR-ABL and TEL-ABL. The differential sensitivity of TEL-ABL and BCR-ABL to asciminib versus ATP-competitive inhibitors may arise from the fact that allosteric inhibition does not completely shut the kinase down, allowing oligomerization to shift the balance of phosphorylation. In contrast, when an ATP-competitive inhibitor is bound it is not possible for the kinase domain to bind ATP and trigger autophosphorylation, leading to more complete suppression of autophosphorylation. A particularly important direction for future work is to study the synergy between ATP-competitive inhibitors and asciminib in TEL-ABL. Although asciminib by itself is ineffective against TEL-ABL, we anticipate that the drug combination will exhibit synergy, through shifting the balance of kinase-phosphatase actions.

Our results demonstrate that the fusion partner can modulate the sensitivity to tyrosine kinase inhibitors. This aligns well with the existing literature on ALK fusions showing that the N-terminal fusion partner influences the sensitivity of ALK to different tyrosine kinase inhibitors (*65*–*68*). Thus, a complete understanding of how a fusion partner affects the properties of the oncogenic protein is essential for the development of new allosteric inhibitors.

## MATERIALS AND METHODS

### Preparation of plasmids

All ABL1 (Uniprot_ID: P00519-1) fusion genes were introduced into the retroviral MSCV-IRES-GFP vector backbone (pMIG, addgene # 20672). p210 BCR-ABL complementary DNA was transferred from the Bcr/Abl P210-pLEF vector (addgene #38158). The TEL-ABL complementary DNA (A form) was generated by fusing TEL exons 1-4 to ABL exon 2. The pMIG vector was digested with EcoRI at a single site to linearize the vector. Amplified DNA fragments were then ligated into the cut pMIG vector using the Gibson assembly protocol. These constructs were used as a template to generate the truncated forms of the two ABL fusion proteins, ABL(core) and ABL(core*Δ*C). These deletion constructs were produced by deleting the ABL1 sequences after residues 581 and 493, respectively. To generate the TEL-ABL A93D and V112E mutants, the TEL sequence was amplified with a pair of oligonucleotide primers designed with mismatching nucleotides at the center of the primers. The mutated TEL DNA fragments and wild type ABL sequence were then ligated into the cut pMIG vector using the Gibson assembly protocol. Chemically competent NEB Stable cells were then transformed with the pMIG vector carrying the ABL fusion gene.

BCR-ABL and TEL-ABL cDNA were cloned into the pEGFP-C1 vector backbone (Clontech, Mountain View, CA), after modifying the vector to contain a biotinylation sequence (Avitag, GLNDIFEAQKIEWHE) (*56*) followed by a flexible linker (GASGASGASGAS) at the N-terminus of monomeric Neon Green protein (mNG). BCR-ABL and TEL-ABL was cloned at the C-terminus of mNG, with a flexible linker sequence (QELSRGGQSG) separating the fluorophore and the coding sequence (i.e., the final construct is organized as avitag-linker-mNG-linker-BCR-ABL/TEL-ABL). To generate TEL-ABL A93D and V112E mutants, the mutated DNA fragment, and the rest of the TEL-ABL sequence were ligated into the modified pEGFP-C1 vector using the Gibson assembly protocol. Chemically competent TOP10 *E.coli* cells were then transformed with the pEGFP-C1 vector carrying the ABL fusion gene. The pET21a-BirA was a gift from Alice Ting (Addgene plasmid # 20857). BirA was cloned into the pSNAPf vector (New England Biolabs, MA) after modifying the vector backbone to remove the SNAP-tag. Each construct was confirmed by DNA sequencing.

### Cell culture

Platinum-E (Plat-E) retroviral packaging cell line was grown at 37°C under 5% CO_2_ in Dulbecco’s Modified Eagle Medium + GlutaMaX (DMEM, Gibco, Thermo Fisher) supplemented with 10% Fetal Bovine Serum (FBS) (Thermo Fisher), 100 U/mL penicillin-streptomycin (Thermo Fisher), 1 μg/mL puromycin (Gibco, Thermo Fisher) and 10 μg/mL blasticidin (InvivoGen). HEK 293T cells were grown at 37°C under 5% CO_2_ in DMEM supplemented with 10% FBS, 50 U/mL penicillin and 50 μg/mL streptomycin. Ba/F3 cells were grown at 37°C under 5% CO_2_ in Roswell Park Memorial Institute 1640 meduim (RPMI, Gibco, Thermo Fisher) supplemented with 10% FBS, 50 U/mL penicillin, 50 μg/mL streptomycin and 2 ng/mL mouse IL-3 (mIL-3, Gibco, Thermo Fisher). 32D cells were maintained in identical culture media with the exception of being supplemented with 15% WEHI-3B cell-conditioned media as a source of IL-3.

### Viral transduction

Plat-E cells were transfected with pMIG-TEL-ABL, pMIG-TEL-ABL(core), pMIG-TEL-ABL(coreΔC), pMIG-BCR-ABL, pMIG-BCR-ABL(core) or pMIG-BCR-ABL(coreΔC) retroviral constructs. Supernatants were collected and filtered through 0.45-μm filters 48 hours after transfection. Ba/F3 cells were incubated 48-72 hours with retroviral medium in the presence of 8 μg/mL polybrene (hexadimethrine bromide, Sigma) and mIL-3 (2 ng/mL). The proportion of transduced Ba/F3 cells (GFP+ population) was determined by flow cytometry, Attune NxT flow cytometer (Thermo Fisher Scientific). mIL-3 was withdrawn, and cells were cultured for 7-10 days without IL-3 for selection of transduced cells. For generation of 32D cell lines, pMIG-TEL-ABL and pMIG-BCR-ABL retroviral constructs were first transfected via Fugene 6 reagent (Promega) and the pEcopak helper plasmid into HEK293T cells. Viral supernatants were harvested at 48 hours and used to transduce 32D cells by spinnoculation, after which cells were resuspended in fresh culture medium and withdrawn from IL-3 for selection as described for Ba/F3 cells above.

### Cellular Proliferation Assay

Ba/F3 cells expressing TEL-ABL or BCR-ABL were plated in 96-well plates and incubated in increasing concentrations of asciminib (Selleckchem), or ponatinib (Selleckchem) for 72 h. Cell viability was assessed by flow cytometry and MTT (3-(4, 5-dimethylthiazolyl-2)-2, 5-diphenyltetrazolium bromide) assay kit (Abcam, ab211091). Data were acquired on an Attune NxT flow cytometer and analyzed with FlowJo v9.9.6 software (TreeStar). IC_50_ values were calculated using Graphpad Prism software using non-linear regression curve fit analysis and are reported as the mean ± SEM of three independent experiments. For 32D cells expressing TEL-ABL or BCR-ABL, similar experiments and analyses were conducted, though cells (1000/well) were distributed into 384-well plates and exposed to the indicated panel of TKIs at concentrations ranging from 0.48 to 2000 nM. Plates were analyzed by MTS-based colorimetric proliferation assay (CellTiter 96 Aqueous One; Promega).

### Flow cytometry analysis

Ba/F3 cells were analyzed in forward scatter (FSC) vs. side scatter (SSC) mode, as well as in fluorescence mode. Regions corresponding to live cells were identified using FSC and SSC, and gates were set to include only these events in the fluorophore intensity distribution and in the subsequent experiments. The distribution of GFP fluorescence for the live-cell population was quantified to determine transduction efficiency.

### Transient transfection

A standard transient-transfection protocol was followed. HEK 293T cells were plated in 6-well plates at a seeding density of 1×10^6^ cells/well one day prior to transfection, to achieve ∼80-90 % confluency at the time of transfection. The Polyethylenimine (PEI) transfection method was used (*69*, *70*). Cells were transfected with a mixture of 2 μg BirA (*56*) and 1 μg of ABL fusion plasmids (full-length BCR-ABL, TEL-ABL or TEL-ABL mutants) at a PEI:DNA ratio of 2.5:1. The cells were then allowed to express the protein for 18–20 hours before harvesting or inhibitor/DMSO treatment.

### Inhibitor treatment in HEK 293T cells

Following an overnight incubation, media including the transfection reagents was removed and replaced by fresh media containing DMSO (less than 0.5%), 0.3 μM ponatinib or asciminib (0.3 μM and 30 μM). After incubation with the DMSO/inhibitor solution for 5 hours, HEK 293T cells were harvested.

### Cell lysis and pulldown of biotinylated ABL fusions in flow chambers

Treated or non-treated HEK 293T cells were harvested and lysed in a lysis buffer containing 25 mM Tris at pH 7.5, 150 mM KCl, 1.5 mM TCEP-HCl, 1% protease-inhibitor cocktail (P8340, Sigma), 0.5% phosphatase-inhibitor cocktail 2 (P5726, Sigma) and 3 (P0044, Sigma), 50 mM NaF, 15 μg/ml benzamidine, 0.1 mM phenylmethanesulfonyl fluoride and 1% NP-40 (Thermo Fisher). The cell lysate was then diluted 100–300 fold in DPBS. Flow chambers were prepared as described previously (*53*). 100 μL of this diluted cell lysate was added to a well in the flow chamber and incubated for 1 min to immobilize the biotinylated mNG-ABL fusion proteins on the surface of the streptavidin-coated glass substrates. The lysate was then washed out with 1 mL of DPBS.

### Immunofluorescence assay with phosphospecific antibodies

The phosphorylation status of the ABL fusion proteins was estimated using a site specific phospho-Tyr 412 antibody (Invitrogen #PA5-99344), or a phospho-Tyr 89 antibody (Cell signaling #3098), or a phospho-Tyr 245 (Invitrogen #44-250), and a pan-phosphotyrosine antibody, 4G10 (Sigma #05-321). The immobilized, treated or untreated ABL fusion proteins were incubated with the phosphotyrosine antibodies using a 1:500 dilution in 5% (w/v) BSA) for 45 min. Subsequently, excess primary antibodies were washed out with 3 mL DPBS. This was followed by a 30 min incubation with an Alexa-labeled secondary antibody that is complementary to the primary antibodies used. Anti-rabbit secondary antibody labeled with Alexa-568 (ThermoFisher Scientific, 1:1000 dilution in 5% BSA) was used for the site-specific primary antibodies (pTyr 412, pTyr 89, and pTyr 245); anti-mouse secondary antibody labeled with Alexa-568 (ThermoFisher Scientific, 1:3000 dilution in 5% BSA) was used for the pTyr 4G10 primary antibody. After incubation with the secondary antibody, the flow chambers were washed again using 3 mL of DPBS and the samples were imaged using Total Internal Reflection Fluorescence (TIRF) microscopy.

### Inhibition assay on a coated glass surface

Following buffer exchange with the gel filtration buffer (25 mM Tris, 150 mM KCl, 1.5 mM TCEP, pH 7.5) to replace DPBS, YopH phosphatase (1 μM in gel filtration buffer) was added to ABL fusion protein captured on the glass. YopH phosphatase was purified as described (*71*). After 30 min, the flow chambers were washed with 2 mL of DPBS. Gel filtration buffer containing 30 nM ponatinib or 30 nM asciminib was then added to the wells. After 30 min of incubation with the inhibitor, wells were washed with 2 mL DPBS and a gel-filtration buffer containing 500 μM ATP and 5 mM MgCl_2_ was added to the wells. The flow chambers were washed with 2 mL DPBS after 30 min. Total phosphorylation and phosphorylation of Tyr 89 were then examined using the immunofluorescence assay described above.

### Single-particle TIRF microscopy

Single-particle TIRF images were acquired on a Nikon Eclipse Ti-inverted microscope equipped with a Nikon 100 × 1.49 numerical aperture oil-immersion TIRF objective, a TIRF illuminator, a Perfect Focus system, and a motorized stage. Images were recorded using an Andor iXon electron-multiplying charge-coupled device camera. The sample was illuminated using the LU-N4 laser unit (Nikon, Tokyo, Japan) with solid state lasers for the 488 nm and 561 nm channels. Lasers were controlled using a built-in acousto-optic tunable filter (AOTF). The 405/488/561/638 nm Quad TIRF filter set (Chroma Technology Corp., Rockingham, Vermont) was used along with supplementary emission filters of 525/50 m, 600/50 m, 700/75 m for 488 nm, 561 nm, 640 nm channel, respectively. 2-color image acquisition was performed by computer-controlled change of illumination and filter sets at 30-35 different positions from an initial reference frame, so as to capture multiple non-overlapping images. Image acquisition was done using the Nikon NIS-Elements software. mNG-ABL fusion proteins, Alexa-568-labeled pan-pTyr and site-specific pTyr 89 antibodies were imaged by illuminating 488 nm laser set to 5.2 mW, and 561 nm laser set to 6.9 mW, respectively. The laser power was measured with the field aperture fully opened. Images were acquired using an exposure time of 50 milliseconds (TEL-ABL) and 75 milliseconds (BCR-ABL) for 488 nm and 80 milliseconds for 561 nm. Epifluorescence images were also acquired using a Nikon Eclipse Ti-inverted microscope with a Nikon 20x objective and an Andor iXon electron-multiplying charge-coupled device camera. A mercury arc lamp (Lumencor Tech., Beaverton, OR) was used for epifluorescence illumination at 488 and 561 nm.

### Analyses of single-particle TIRF data

Individual single particles were detected in the green and red channels and localized using the single-particle tracking plugin TrackMate in ImageJ (*72*, *73*). The particles were localized with the Difference of Gaussian (DoG) detector with an initial diameter set to five pixels. The detection threshold value in TrackMate is set to 60 (mNG-TEL-ABL at 488nm), 80 (mNG-BCR-ABL at 488nm) or 50 (phospho antibodies at 561nm) depending on the fusion protein and the wavelength at which the image is acquired. The particles outside the central area of 350 × 350 square pixels were excluded due to heterogeneous TIRF illumination. No further filtering processes were applied in TrackMate.

The distribution of fluorescence intensities for the phospho-antibodies, and the fraction of ABL fusion proteins showing any detectable phosphorylation were analyzed using custom in-house software written in MATLAB (these programs are provided as a source code file). For analyzing the fraction of spots that show any detectable phosphorylation at a given phosphosite, the XY-coordinates for each particle were extracted from the TrackMate data. For each mNG-ABL fusion protein, distances to all the antibody particles were computed, and colocalization was identified if the inter-particle distance was below two pixels. The fraction of mNG-ABL fusion proteins with detectable phosphorylation was computed as a ratio of the number of ABL fusion protein colocalized with antibody to the total number of ABL fusion complexes. In a typical case, false-positive colocalization due to coincidence was less than 0.5%, as determined from analyzing multiple, unrelated sets of images with comparable particle density. The intensity values acquired from TrackMate statistics data for the pan-pTyr and pTyr 89 antibody channels were used to calculate an intensity histogram, for the colocalized (i.e., corresponding to individual complexes that show detectable phosphorylation) and the non-colocalized spots (i.e., corresponding to individual complexes that do not show any detectable phosphorylation with intensity values in the phosphospecific antibody channel being zero). This results in scaling of the area under the histogram for the antibody intensity by a factor that takes into account the fraction of holoenzymes that show no detectable phosphorylation. A bin width of 500 was used for computing the histogram. After calculating the distribution, we plotted the intensity histogram for only the colocalized spots and an intensity cutoff of about 50000 arbitrary units. For the analysis of the mNG step-photobleaching data, the photobleaching traces for each spot were built by plotting the maximum intensities of mNG-ABL fusion proteins as a function of time with MATLAB. The number of single photobleaching events were counted manually by inspecting the photobleaching trace for every spot. Single-step, two-step and three-step bleaching traces were clearly identified, but multistep photobleaching traces exhibited relatively unclear, distinct bleaching events. A similar analysis was used for mNG-TEL-ABL mutants, which exhibited mostly single-step traces.

### Analysis of oligomeric states of ABL fusion proteins from single-particle TIRF images

The fluorescence intensity distributions for the fluorescent protein tags showed obvious peaks corresponding to different numbers of fluorescent molecules in immobilized complexes (Supp Figure 6A). This motivated us to carry out a quantitative analysis of the intensity distributions, with the aim of inferring the distribution of oligomer sizes.

First, we approximate the point spread function (PSF) by a circularly-symmetric Gaussian to estimate the intensity from multiple pixels and prevent varying defocus from being reflected in the intensities. We note that the true PSF of a rotationally-averaged point dipole in TIRF is not Gaussian (*74*), but the Gaussian approximation is very accurate (*75*). Specifically, we use the Laplacian of Gaussian, as implemented in scikit-image (*76*), to identify particle locations and optimize the Gaussian scale per-particle, then quantify each particle as the intensity of the image convolved with a Gaussian of that scale evaluated at that pixel. For monomeric mutants, there is a plateau in the number of blobs identified at a given threshold. This occurs in the range of thresholds between excluding mostly background locations and excluding mostly immobilized fluorophores from the identified particles. We chose a threshold for particle identification that is near this local minimum in the number of particles newly excluded as the threshold increased.

We also applied a vignetting correction to account for uneven illumination and imaging across the frame. Here, we used a Gaussian profile to describe the decay from the center of vignetting to the corners. Combined with a lognormal intensity distribution for the background, our model becomes:

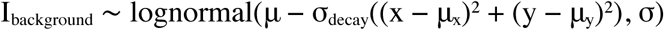

In both dimensions, our maximum likelihood estimate of the vignetting center is within 10% of the frame center, with σ_decay_ ≈ 1000 pixels corresponding to a maximum 15% correction in the corner of the 512×512 frame.

Images for each sample were collected at multiple locations imaged consecutively in the same flowcell. While there is no evidence of severe photobleaching of immobilized fluorophores over consecutive exposures, the background intensity drops noticeably over the first few frames. For this reason, the background intensity was estimated and subtracted per-image. After applying the invariant vignetting correction and identifying particles using an invariant threshold, pixels 4σ from any particles were used for maximum likelihood estimation of a lognormal background distribution, then the mode, e^μ−σ2^, was subtracted.

At this point, it was obvious from the intensity histograms of the four samples that the two TEL-ABL mutants (TEL-ABL A93D and TEL-ABL V112E) appeared as a single, presumably monomeric, species (Supp Figure 6B). An identical peak was present in BCR-ABL and TEL-ABL wild-types. While the only other major peak of BCR-ABL seemed dimeric, TEL-ABL exhibited a broad distribution of higher intensities in addition to the single-fluorophore peak. The strong information about monomer and dimer intensities from the mutant and BCR-ABL samples provides a good foundation from which to deconvolve the intensity distibutions to obtain approximate oligomeric state distributions.

Here, we assume binomial-lognormal mixture models of the form:

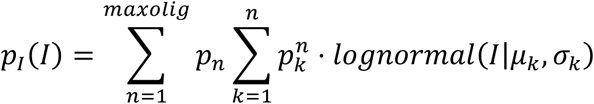

That is, intensities are lognormally-distributed for a given number of bright fluorophores (*77*). The number of bright fluorophores given a particular oligomeric state is binomially-distributed, with a probability shared across all samples, and the zero-bright population unobserved:

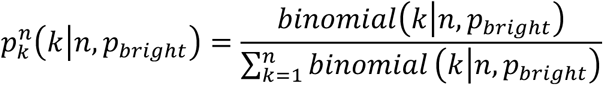

We modeled lognormal parameters for multiple bright fluorophores with three different assumptions or approximations, but no free parameters:

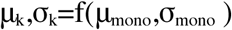

The simplest model results from multiplicative combination of the individual fluorophore intensities (*77*) or alternatively, equally-spaced *e*^*μ_k_*^ with *σ*_*k*_ constant:

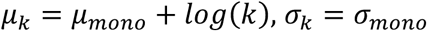

The distribution of a sum of lognormally-distributed random variables, as would be expected for fluorophores within a complex contributed independently to overall intensity, can be approximated by a lognormal with parameters chosen to match the first two moments:

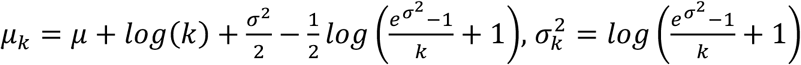

We can as well include an explicit background distribution in the sum of lognormals:

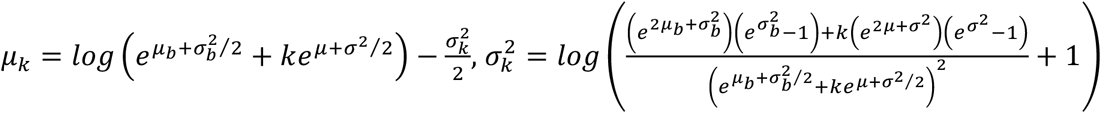

A numerically- and computationally-attractive approximation to the moment-matched parameters without background distribution, exact for *k* = 1 and with indistinguishable density for this data is:

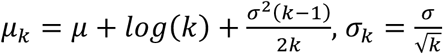

We assume that the TEL-ABL mutants are monomeric, but make no assumptions and place no priors on the distributions of BCR-ABL and TEL-ABL wild-type beyond truncating at 10-mers. We sampled from these models given all four experimental samples simultaneously using CmdStanPy (*78*) and present the intensity distribution predicted by the mean parameter values, and the mean and samples for the oligomeric state distributions (*79*–*81*). All three models give qualitatively similar results for likely oligomeric distributions of TEL-ABL.

A Jupyter notebook containing the full data analysis is available at: https://github.com/kentgorday/kuriyanlab_python-workshops/blob/master/bonus/inferring_lognormal-mixture_demo.ipynb

## Author contributions

Conceptualization: SM, JK

Resources: SM, CAE, KG, JTG, JK

Data Curation: SM, KG, ES, WZ

Software: SM, KG, ES, WZ

Methodology: SM, CAE, KG, JTG, BJD, JK

Investigation: SM, CAE, KG, ES, WZ, JK

Visualization: SM, JK

Funding acquisition: JK

Project administration: JK

Supervision: JK

Writing – original draft: SM, JK

Writing – review & editing: SM, CAE, KG, JTG, BJD, JK

## Data availability

All data generated or analyzed during this study are summarized in the manuscript, figures, and supplementary files. The in-house Matlab programs that are used for colocalization, intensity, and photobleaching analysis will be provided as Source code files. A Jupyter notebook containing the analysis of oligomeric states of ABL fusions is available at https://github.com/kentgorday/kuriyanlab_python-workshops/blob/master/bonus/inferring_lognormal-mixture_demo.ipynb

## References and Notes

1. O. Hantschel, Structure, regulation, signaling, and targeting of abl kinases in cancer. Genes Cancer. 3, 436–446 (2012).

2. E. Shtivelman, B. Lifshitz, R. P. Gale, E. Canaani, Fused transcript of abl and bcr genes in chronic myelogenous leukaemia. Nature. 315, 550–554 (1985).

3. B. D. Lichty, A. Keating, J. Callum, K. Yee, R. Croxford, G. Corpus, B. Nwachukwu, P. Kim, J. Guo, S. Kamel-Reid, Expression of p210 and p190 BCR-ABL due to alternative splicing in chronic myelogenous leukaemia. Br. J. Haematol. 103, 711– 715 (1998).

4. V. Pullarkat, M. L. Slovak, K. J. Kopecky, S. J. Forman, F. R. Appelbaum, Impact of cytogenetics on the outcome of adult acute lymphoblastic leukemia: results of Southwest Oncology Group 9400 study. Blood. 111, 2563–2572 (2008).

5. P. Papadopoulos, S. A. Ridge, C. A. Boucher, C. Stocking, L. M. Wiedemann, The novel activation of ABL by fusion to an ets-related gene, TEL. Cancer Res. 55, 34–38 (1995).

6. H. Van Limbergen, H. B. Beverloo, E. van Drunen, A. Janssens, K. Hählen, B. Poppe, N. Van Roy, P. Marynen, A. De Paepe, R. Slater, F. Speleman, Molecular cytogenetic and clinical findings in ETV6/ABL1-positive leukemia. Genes Chromosomes Cancer. 30, 274–282 (2001).

7. T. R. Golub, A. Goga, G. F. Barker, D. E. Afar, J. McLaughlin, S. K. Bohlander, J. D. Rowley, O. N. Witte, D. G. Gilliland, Oligomerization of the ABL tyrosine kinase by the Ets protein TEL in human leukemia. Mol. Cell. Biol. 16, 4107–4116 (1996).

8. T. Burjanivova, J. Madzo, K. Muzikova, C. Meyer, B. Schneider, F. Votava, R. Marschalek, J. Stary, J. Trka, J. Zuna, Prenatal origin of childhood AML occurs less frequently than in childhood ALL. BMC Cancer. 6, 100 (2006).

9. R. La Starza, M. Trubia, N. Testoni, E. Ottaviani, E. Belloni, B. Crescenzi, M. Martelli, G. Flandrin, P. G. Pelicci, C. Mecucci, Clonal eosinophils are a morphologic hallmark of ETV6/ABL1 positive acute myeloid leukemia. Haematologica. 87, 789– 794 (2002).

10. H. Lin, J. Q. Guo, M. Andreeff, R. B. Arlinghaus, Detection of dual TEL-ABL transcripts and a Tel-Abl protein containing phosphotyrosine in a chronic myeloid leukemia patient. Leukemia. 16, 294–297 (2002).

11. S. G. O’Brien, S. A. D. Vieira, S. Connors, N. Bown, J. Chang, R. Capdeville, J. V. Melo, Transient response to imatinib mesylate (STI571) in a patient with the ETV6-ABL t(9;12) translocation. Blood. 99, 3465–3467 (2002).

12. S. Meyer-Monard, D. Mühlematter, A. Streit, A. J. Chase, A. Gratwohl, N. C. P. Cross, M. Jotterand, A. Tichelli, Broad molecular screening of an unclassifiable myeloproliferative disorder reveals an unexpected ETV6/ABL1 fusion transcript. Leukemia. 19, 1096–1099 (2005).

13. C. A. Tirado, S. Sebastian, J. O. Moore, J. Z. Gong, B. K. Goodman, Molecular and cytogenetic characterization of a novel rearrangement involving chromosomes 9, 12, and 17 resulting in ETV6 (TEL) and ABL fusion. Cancer Genet. Cytogenet. 157, 74–77 (2005).

14. M.-J. Mozziconacci, D. Sainty, C. Chabannon, A fifteen-year cytogenetic remission following interferon treatment in a patient with an indolent ETV6-ABL positive myeloproliferative syndrome. Am. J. Hematol. 82, 688–689 (2007).

15. J. Baeumler, K. Szuhai, J. H. F. Falkenburg, M. L. J. van Schie, O. G. Ottmann, B. A. Nijmeijer, Establishment and cytogenetic characterization of a human acute lymphoblastic leukemia cell line (ALL-VG) with ETV6/ABL1 rearrangement. Cancer Genet. Cytogenet. 185, 37–42 (2008).

16. N. Kawamata, A. Dashti, D. Lu, B. Miller, H. P. Koeffler, R. Schreck, S. Moore, S. Ogawa, Chronic phase of ETV6-ABL1 positive CML responds to imatinib. Genes Chromosomes Cancer. 47, 919–921 (2008).

17. J. C. Kelly, N. Shahbazi, J. Scheerle, J. Jahn, S. Suchen, N. C. Christacos, P. N. Mowrey, M. H. Witt, A. Hostetter, A. M. Meloni-Ehrig, Insertion (12;9)(p13;q34q34): a cryptic rearrangement involving ABL1/ETV6 fusion in a patient with Philadelphia-negative chronic myeloid leukemia. Cancer Genet. Cytogenet. 192, 36–39 (2009).

18. R. Nand, C. Bryke, S. H. Kroft, A. Divgi, C. Bredeson, E. Atallah, Myeloproliferative disorder with eosinophilia and ETV6-ABL gene rearrangement: efficacy of second-generation tyrosine kinase inhibitors. Leuk. Res. 33, 1144–1146 (2009).

19. H. Pluk, K. Dorey, G. Superti-Furga, Autoinhibition of c-Abl. Cell. 108, 247–259 (2002).

20. O. Hantschel, B. Nagar, S. Guettler, J. Kretzschmar, K. Dorey, J. Kuriyan, G. Superti-Furga, A myristoyl/phosphotyrosine switch regulates c-Abl. Cell. 112, 845–857 (2003).

21. B. Nagar, O. Hantschel, M. Seeliger, J. M. Davies, W. I. Weis, G. Superti-Furga, J. Kuriyan, Organization of the SH3-SH2 unit in active and inactive forms of the c-Abl tyrosine kinase. Mol. Cell. 21, 787–798 (2006).

22. B. Nagar, O. Hantschel, M. A. Young, K. Scheffzek, D. Veach, W. Bornmann, B. Clarkson, G. Superti-Furga, J. Kuriyan, Structural basis for the autoinhibition of c-Abl tyrosine kinase. Cell. 112, 859–871 (2003).

23. Y. He, J. A. Wertheim, L. Xu, J. P. Miller, F. G. Karnell, J. K. Choi, R. Ren, W. S. Pear, The coiled-coil domain and Tyr177 of bcr are required to induce a murine chronic myelogenous leukemia-like disease by bcr/abl. Blood. 99, 2957–2968 (2002).

24. X. Zhang, R. Subrahmanyam, R. Wong, A. W. Gross, R. Ren, The NH(2)-terminal coiled-coil domain and tyrosine 177 play important roles in induction of a myeloproliferative disease in mice by Bcr-Abl. Mol. Cell. Biol. 21, 840–853 (2001).

25. J. R. McWhirter, D. L. Galasso, J. Y. Wang, A coiled-coil oligomerization domain of Bcr is essential for the transforming function of Bcr-Abl oncoproteins. Mol. Cell. Biol. 13, 7587–7595 (1993).

26. C. M. Taylor, A. E. Keating, Orientation and oligomerization specificity of the Bcr coiled-coil oligomerization domain. Biochemistry. 44, 16246–16256 (2005).

27. X. Zhao, S. Ghaffari, H. Lodish, V. N. Malashkevich, P. S. Kim, Structure of the Bcr-Abl oncoprotein oligomerization domain. Nat. Struct. Biol. 9, 117–120 (2002).

28. H. H. Tran, C. A. Kim, S. Faham, M.-C. Siddall, J. U. Bowie, Native interface of the SAM domain polymer of TEL. BMC Struct. Biol. 2, 5 (2002).

29. C. A. Kim, M. L. Phillips, W. Kim, M. Gingery, H. H. Tran, M. A. Robinson, S. Faham, J. U. Bowie, Polymerization of the SAM domain of TEL in leukemogenesis and transcriptional repression. EMBO J. 20, 4173–4182 (2001).

30. B. J. Druker, S. Tamura, E. Buchdunger, S. Ohno, G. M. Segal, S. Fanning, J. Zimmermann, N. B. Lydon, Effects of a selective inhibitor of the Abl tyrosine kinase on the growth of Bcr-Abl positive cells. Nat. Med. 2, 561–566 (1996).

31. J. S. Tokarski, J. A. Newitt, C. Y. J. Chang, J. D. Cheng, M. Wittekind, S. E. Kiefer, K. Kish, F. Y. F. Lee, R. Borzillerri, L. J. Lombardo, D. Xie, Y. Zhang, H. E. Klei, The structure of Dasatinib (BMS-354825) bound to activated ABL kinase domain elucidates its inhibitory activity against imatinib-resistant ABL mutants. Cancer Res. 66, 5790–5797 (2006).

32. L. J. Lombardo, F. Y. Lee, P. Chen, D. Norris, J. C. Barrish, K. Behnia, S. Castaneda, L. A. M. Cornelius, J. Das, A. M. Doweyko, C. Fairchild, J. T. Hunt, I. Inigo, K. Johnston, A. Kamath, D. Kan, H. Klei, P. Marathe, S. Pang, R. Peterson, S. Pitt, G. L. Schieven, R. J. Schmidt, J. Tokarski, M.-L. Wen, J. Wityak, R. M. Borzilleri, Discovery of N-(2-chloro-6-methyl-phenyl)-2-(6-(4-(2-hydroxyethyl)-piperazin-1-yl)-2-methylpyrimidin-4- ylamino)thiazole-5-carboxamide (BMS-354825), a dual Src/Abl kinase inhibitor with potent antitumor activity in preclinical assays. J. Med. Chem. 47, 6658–6661 (2004).

33. E. Weisberg, P. W. Manley, W. Breitenstein, J. Brüggen, S. W. Cowan-Jacob, A. Ray, B. Huntly, D. Fabbro, G. Fendrich, E. Hall-Meyers, A. L. Kung, J. Mestan, G. Q. Daley, L. Callahan, L. Catley, C. Cavazza, M. Azam, D. Neuberg, R. D. Wright, D. G. Gilliland, J. D. Griffin, Characterization of AMN107, a selective inhibitor of native and mutant Bcr-Abl. Cancer Cell. 7, 129–141 (2005).

34. T. O’Hare, W. C. Shakespeare, X. Zhu, C. A. Eide, V. M. Rivera, F. Wang, L. T. Adrian, T. Zhou, W.-S. Huang, Q. Xu, C. A. Metcalf, J. W. Tyner, M. M. Loriaux, A. S. Corbin, S. Wardwell, Y. Ning, J. A. Keats, Y. Wang, R. Sundaramoorthi, M. Thomas, D. Zhou, J. Snodgrass, L. Commodore, T. K. Sawyer, D. C. Dalgarno, M. W. N. Deininger, B. J. Druker, T. Clackson, AP24534, a pan-BCR-ABL inhibitor for chronic myeloid leukemia, potently inhibits the T315I mutant and overcomes mutation-based resistance. Cancer Cell. 16, 401–412 (2009).

35. J. Schoepfer, W. Jahnke, G. Berellini, S. Buonamici, S. Cotesta, S. W. Cowan-Jacob, S. Dodd, P. Drueckes, D. Fabbro, T. Gabriel, J.-M. Groell, R. M. Grotzfeld, A. Q. Hassan, C. Henry, V. Iyer, D. Jones, F. Lombardo, A. Loo, P. W. Manley, X. Pellé, G. Rummel, B. Salem, M. Warmuth, A. A. Wylie, T. Zoller, A. L. Marzinzik, P. Furet, Discovery of Asciminib (ABL001), an Allosteric Inhibitor of the Tyrosine Kinase Activity of BCR-ABL1. J. Med. Chem. 61, 8120–8135 (2018).

36. A. A. Wylie, J. Schoepfer, W. Jahnke, S. W. Cowan-Jacob, A. Loo, P. Furet, A. L. Marzinzik, X. Pelle, J. Donovan, W. Zhu, S. Buonamici, A. Q. Hassan, F. Lombardo, V. Iyer, M. Palmer, G. Berellini, S. Dodd, S. Thohan, H. Bitter, S. Branford, D. M. Ross, T. P. Hughes, L. Petruzzelli, K. G. Vanasse, M. Warmuth, F. Hofmann, N. J. Keen, W. R. Sellers, The allosteric inhibitor ABL001 enables dual targeting of BCR-ABL1. Nature. 543, 733–737 (2017).

37. C. A. Eide, M. S. Zabriskie, S. L. Savage Stevens, O. Antelope, N. A. Vellore, H. Than, A. R. Schultz, P. Clair, A. D. Bowler, A. D. Pomicter, D. Yan, A. V. Senina, W. Qiang, T. W. Kelley, P. Szankasi, M. C. Heinrich, J. W. Tyner, D. Rea, J.-M. Cayuela, D.-W. Kim, C. E. Tognon, T. O’Hare, B. J. Druker, M. W. Deininger, Combining the Allosteric Inhibitor Asciminib with Ponatinib Suppresses Emergence of and Restores Efficacy against Highly Resistant BCR-ABL1 Mutants. Cancer Cell. 36, 431–443.e5 (2019).

38. N. Curik, A. Laznicka, V. Polivkova, J. Krizkova, E. Pokorna, P. Semerak, P. Suchankova, P. Burda, A. Hochhaus, K. Machova Polakova, Combination therapies with ponatinib and asciminib in a preclinical model of chronic myeloid leukemia blast crisis with compound mutations. Leukemia. 38, 1415–1418 (2024).

39. J. Cortes, F. Lang, D. Rea, A. Hochhaus, M. Breccia, Y. T. Goh, M. C. Heinrich, T. P. Hughes, J. J. W. M. Janssen, P. le Coutre, H. Minami, K. Sasaki, D. J. DeAngelo, G. Sanchez-Olle, N. Pognan, J. Jose, M. Hoch, M. Mauro, Asciminib (ASC) in Combination with Imatinib (IMA), Nilotinib (NIL), or Dasatinib (DAS) May be a Potential Treatment (Tx) Option in Patients (Pts) with Philadelphia Chromosome-Positive Chronic Myeloid Leukemia in Chronic Phase or Accelerated Phase (Ph+ CML-CP/AP): Final Results from the Asciminib Phase 1 Study. Blood. 142, 868–868 (2023).

40. B. Oruganti, E. Lindahl, J. Yang, W. Amiri, R. Rahimullah, R. Friedman, Allosteric enhancement of the BCR-Abl1 kinase inhibition activity of nilotinib by cobinding of asciminib. J. Biol. Chem. 298, 102238 (2022).

41. N. Okamoto, K. Yagi, S. Imawaka, M. Takaoka, F. Aizawa, T. Niimura, M. Goda, K. Miyata, K. Kawada, Y. Izawa-Ishizawa, S. Sakaguchi, K. Ishizawa, Asciminib, a novel allosteric inhibitor ofBCR-ABL1, shows synergistic effects when used in combination with imatinib with or without drug resistance. Pharmacol. Res. Perspect. 12 (2024), doi:10.1002/prp2.1214.

42. F. J. Adrián, Q. Ding, T. Sim, A. Velentza, C. Sloan, Y. Liu, G. Zhang, W. Hur, S. Ding, P. Manley, J. Mestan, D. Fabbro, N. S. Gray, Allosteric inhibitors of Bcr-abl-dependent cell proliferation. Nat. Chem. Biol. 2, 95–102 (2006).

43. D. Fabbro, P. W. Manley, W. Jahnke, J. Liebetanz, A. Szyttenholm, G. Fendrich, A. Strauss, J. Zhang, N. S. Gray, F. Adrian, M. Warmuth, X. Pelle, R. Grotzfeld, F. Berst, A. Marzinzik, S. W. Cowan-Jacob, P. Furet, J. Mestan, Inhibitors of the Abl kinase directed at either the ATP- or myristate-binding site. Biochim. Biophys. Acta. 1804, 454–462 (2010).

44. J. Zhang, F. J. Adrián, W. Jahnke, S. W. Cowan-Jacob, A. G. Li, R. E. Iacob, T. Sim, J. Powers, C. Dierks, F. Sun, G.-R. Guo, Q. Ding, B. Okram, Y. Choi, A. Wojciechowski, X. Deng, G. Liu, G. Fendrich, A. Strauss, N. Vajpai, S. Grzesiek, T. Tuntland, Y. Liu, B. Bursulaya, M. Azam, P. W. Manley, J. R. Engen, G. Q. Daley, M. Warmuth, N. S. Gray, Targeting Bcr-Abl by combining allosteric with ATP-binding-site inhibitors. Nature. 463, 501–506 (2010).

45. T. P. Hughes, M. J. Mauro, J. E. Cortes, H. Minami, D. Rea, D. J. DeAngelo, M. Breccia, Y.-T. Goh, M. Talpaz, A. Hochhaus, P. le Coutre, O. Ottmann, M. C. Heinrich, J. L. Steegmann, M. W. N. Deininger, J. J. W. M. Janssen, F.-X. Mahon, Y. Minami, D. Yeung, D. M. Ross, M. S. Tallman, J. H. Park, B. J. Druker, D. Hynds, Y. Duan, C. Meille, F. Hourcade-Potelleret, K. G. Vanasse, F. Lang, D.-W. Kim, Asciminib in Chronic Myeloid Leukemia after ABL Kinase Inhibitor Failure. N. Engl. J. Med. 381, 2315–2326 (2019).

46. P. W. Manley, L. Barys, S. W. Cowan-Jacob, The specificity of asciminib, a potential treatment for chronic myeloid leukemia, as a myristate-pocket binding ABL inhibitor and analysis of its interactions with mutant forms of BCR-ABL1 kinase. Leuk. Res. 98, 106458 (2020).

47. S. Chen, L. P. O’Reilly, T. E. Smithgall, J. R. Engen, Tyrosine phosphorylation in the SH3 domain disrupts negative regulatory interactions within the c-Abl kinase core. J. Mol. Biol. 383, 414–423 (2008).

48. B. B. Brasher, R. A. Van Etten, c-Abl has high intrinsic tyrosine kinase activity that is stimulated by mutation of the Src homology 3 domain and by autophosphorylation at two distinct regulatory tyrosines. J. Biol. Chem. 275, 35631–35637 (2000).

49. G. Q. Daley, D. Baltimore, Transformation of an interleukin 3-dependent hematopoietic cell line by the chronic myelogenous leukemia-specific P210bcr/abl protein. Proc Natl Acad Sci USA. 85, 9312–9316 (1988).

50. J. S. Greenberger, M. A. Sakakeeny, R. K. Humphries, C. J. Eaves, R. J. Eckner, Demonstration of permanent factor-dependent multipotential (erythroid/neutrophil/basophil) hematopoietic progenitor cell lines. Proc Natl Acad Sci USA. 80, 2931–2935 (1983).

51. M. Valtieri, D. J. Tweardy, D. Caracciolo, K. Johnson, F. Mavilio, S. Altmann, D. Santoli, G. Rovera, Cytokine-dependent granulocytic differentiation. Regulation of proliferative and differentiative responses in a murine progenitor cell line. J. Immunol. 138, 3829–3835 (1987).

52. K. J. Johnson, I. J. Griswold, T. O’Hare, A. S. Corbin, M. Loriaux, M. W. Deininger, B. J. Druker, A BCR-ABL mutant lacking direct binding sites for the GRB2, CBL and CRKL adapter proteins fails to induce leukemia in mice. PLoS ONE. 4, e7439 (2009).

53. M. Bhattacharyya, Y. K. Lee, S. Muratcioglu, B. Qiu, P. Nyayapati, H. Schulman, J. T. Groves, J. Kuriyan, Flexible linkers in CaMKII control the balance between activating and inhibitory autophosphorylation. eLife. 9 (2020), doi:10.7554/eLife.53670.

54. K. N. Fish, Curr. Protoc. Cytom., in press, doi:10.1002/0471142956.cy1218s50.

55. N. C. Shaner, G. G. Lambert, A. Chammas, Y. Ni, P. J. Cranfill, M. A. Baird, B. R. Sell, J. R. Allen, R. N. Day, M. Israelsson, M. W. Davidson, J. Wang, A bright monomeric green fluorescent protein derived from Branchiostoma lanceolatum. Nat. Methods. 10, 407–409 (2013).

56. D. Beckett, E. Kovaleva, P. J. Schatz, A minimal peptide substrate in biotin holoenzyme synthetase-catalyzed biotinylation. Protein Sci. 8, 921–929 (1999).

57. M. A. Meyn, M. B. Wilson, F. A. Abdi, N. Fahey, A. P. Schiavone, J. Wu, J. M. Hochrein, J. R. Engen, T. E. Smithgall, Src family kinases phosphorylate the Bcr-Abl SH3-SH2 region and modulate Bcr-Abl transforming activity. J. Biol. Chem. 281, 30907–30916 (2006).

58. J. B. Bliska, K. L. Guan, J. E. Dixon, S. Falkow, Tyrosine phosphate hydrolysis of host proteins by an essential Yersinia virulence determinant. Proc Natl Acad Sci USA. 88, 1187–1191 (1991).

59. D. Réa, T. P. Hughes, Development of asciminib, a novel allosteric inhibitor of BCR-ABL1. Crit. Rev. Oncol. Hematol. 171, 103580 (2022).

60. I. B. Leske, O. Hantschel, The e13a3 (b2a3) and e14a3 (b3a3) BCR::ABL1 isoforms are resistant to asciminib. Leukemia (2024), doi:10.1038/s41375-024-02314-7.

61. L. Skora, J. Mestan, D. Fabbro, W. Jahnke, S. Grzesiek, NMR reveals the allosteric opening and closing of Abelson tyrosine kinase by ATP-site and myristoyl pocket inhibitors. Proc Natl Acad Sci USA. 110, E4437–45 (2013).

62. R. Sonti, I. Hertel-Hering, A. J. Lamontanara, O. Hantschel, S. Grzesiek, ATP site ligands determine the assembly state of the abelson kinase regulatory core via the activation loop conformation. J. Am. Chem. Soc. 140, 1863–1869 (2018).

63. C. Kim, H. Ludewig, A. Hadzipasic, S. Kutter, V. Nguyen, D. Kern, A biophysical framework for double-drugging kinases. Proc Natl Acad Sci USA. 120, e2304611120 (2023).

64. T. K. Johnson, D. A. Bochar, N. M. Vandecan, J. Furtado, M. P. Agius, S. Phadke, M. B. Soellner, Synergy and Antagonism between Allosteric and Active-Site Inhibitors of Abl Tyrosine Kinase. Angew. Chem. Int. Ed. 60, 20196–20199 (2021).

65. C. G. Woo, S. Seo, S. W. Kim, S. J. Jang, K. S. Park, J. Y. Song, B. Lee, M. W. Richards, R. Bayliss, D. H. Lee, J. Choi, Differential protein stability and clinical responses of EML4-ALK fusion variants to various ALK inhibitors in advanced ALK-rearranged non-small cell lung cancer. Ann. Oncol. 28, 791–797 (2017).

66. J. J. Lin, V. W. Zhu, S. Yoda, B. Y. Yeap, A. B. Schrock, I. Dagogo-Jack, N. A. Jessop, G. Y. Jiang, L. P. Le, K. Gowen, P. J. Stephens, J. S. Ross, S. M. Ali, V. A. Miller, M. L. Johnson, C. M. Lovly, A. N. Hata, J. F. Gainor, A. J. Iafrate, A. T. Shaw, S.-H. I. Ou, Impact of EML4-ALK Variant on Resistance Mechanisms and Clinical Outcomes in ALK-Positive Lung Cancer. J. Clin. Oncol. 36, 1199–1206 (2018).

67. T. Yoshida, Y. Oya, K. Tanaka, J. Shimizu, Y. Horio, H. Kuroda, Y. Sakao, T. Hida, Y. Yatabe, Differential Crizotinib Response Duration Among ALK Fusion Variants in ALK-Positive Non-Small-Cell Lung Cancer. J. Clin. Oncol. 34, 3383–3389 (2016).

68. M. A. Childress, S. M. Himmelberg, H. Chen, W. Deng, M. A. Davies, C. M. Lovly, ALK fusion partners impact response to ALK inhibition: differential effects on sensitivity, cellular phenotypes, and biochemical properties. Mol. Cancer Res. 16, 1724–1736 (2018).

69. O. Boussif, F. Lezoualc’h, M. A. Zanta, M. D. Mergny, D. Scherman, B. Demeneix, J. P. Behr, A versatile vector for gene and oligonucleotide transfer into cells in culture and in vivo: polyethylenimine. Proc Natl Acad Sci USA. 92, 7297–7301 (1995).

70. N. D. Sonawane, F. C. Szoka, A. S. Verkman, Chloride accumulation and swelling in endosomes enhances DNA transfer by polyamine-DNA polyplexes. J. Biol. Chem. 278, 44826–44831 (2003).

71. Z. Y. Zhang, J. C. Clemens, H. L. Schubert, J. A. Stuckey, M. W. Fischer, D. M. Hume, M. A. Saper, J. E. Dixon, Expression, purification, and physicochemical characterization of a recombinant Yersinia protein tyrosine phosphatase. J. Biol. Chem. 267, 23759–23766 (1992).

72. K. Jaqaman, D. Loerke, M. Mettlen, H. Kuwata, S. Grinstein, S. L. Schmid, G. Danuser, Robust single-particle tracking in live-cell time-lapse sequences. Nat. Methods. 5, 695–702 (2008).

73. J. Schindelin, I. Arganda-Carreras, E. Frise, V. Kaynig, M. Longair, T. Pietzsch, S. Preibisch, C. Rueden, S. Saalfeld, B. Schmid, J.-Y. Tinevez, D. J. White, V. Hartenstein, K. Eliceiri, P. Tomancak, A. Cardona, Fiji: an open-source platform for biological-image analysis. Nat. Methods. 9, 676–682 (2012).

74. K. I. Mortensen, L. S. Churchman, J. A. Spudich, H. Flyvbjerg, Optimized localization analysis for single-molecule tracking and super-resolution microscopy. Nat. Methods. 7, 377–381 (2010).

75. K. I. Mortensen, H. Flyvbjerg, “Calibration-on-the-spot”: How to calibrate an EMCCD camera from its images. Sci. Rep. 6, 28680 (2016).

76. S. van der Walt, J. L. Schönberger, J. Nunez-Iglesias, F. Boulogne, J. D. Warner, N. Yager, E. Gouillart, T. Yu, scikit-image contributors, scikit-image: image processing in Python. PeerJ. 2, e453 (2014).

77. S. A. Mutch, B. S. Fujimoto, C. L. Kuyper, J. S. Kuo, S. M. Bajjalieh, D. T. Chiu, Deconvolving single-molecule intensity distributions for quantitative microscopy measurements. Biophys. J. 92, 2926–2943 (2007).

78. B. Carpenter, A. Gelman, M. D. Hoffman, D. Lee, B. Goodrich, M. Betancourt, M. A. Brubaker, J. Guo, P. Li, A. Riddell, Stan: A probabilistic programming language. J. Stat. Softw. 76 (2017), doi:10.18637/jss.v076.i01.

79. C. R. Harris, K. J. Millman, S. J. van der Walt, R. Gommers, P. Virtanen, D. Cournapeau, E. Wieser, J. Taylor, S. Berg, N. J. Smith, R. Kern, M. Picus, S. Hoyer, M. H. van Kerkwijk, M. Brett, A. Haldane, J. F. Del Río, M. Wiebe, P. Peterson, P. Gérard-Marchant, K. Sheppard, T. Reddy, W. Weckesser, H. Abbasi, C. Gohlke, T. E. Oliphant, Array programming with NumPy. Nature. 585, 357–362 (2020).

80. P. Virtanen, R. Gommers, T. E. Oliphant, M. Haberland, T. Reddy, D. Cournapeau, E. Burovski, P. Peterson, W. Weckesser, J. Bright, S. J. van der Walt, M. Brett, J. Wilson, K. J. Millman, N. Mayorov, A. R. J. Nelson, E. Jones, R. Kern, E. Larson, C. J. Carey, İ. Polat, Y. Feng, E. W. Moore, J. VanderPlas, D. Laxalde, J. Perktold, R. Cimrman, I. Henriksen, E. A. Quintero, C. R. Harris, A. M. Archibald, A. H. Ribeiro, F. Pedregosa, P. van Mulbregt, SciPy 1.0 Contributors, SciPy 1.0: fundamental algorithms for scientific computing in Python. Nat. Methods. 17, 261–272 (2020).

81. J. D. Hunter, Matplotlib: A 2D Graphics Environment. Comput. Sci. Eng. 9, 90–95 (2007).

