## Supplemental Figure 1 for "Autophosphorylation of the oncogenic protein TEL-ABL confers resistance to the allosteric ABL inhibitor asciminib"

### Supplementary Figures

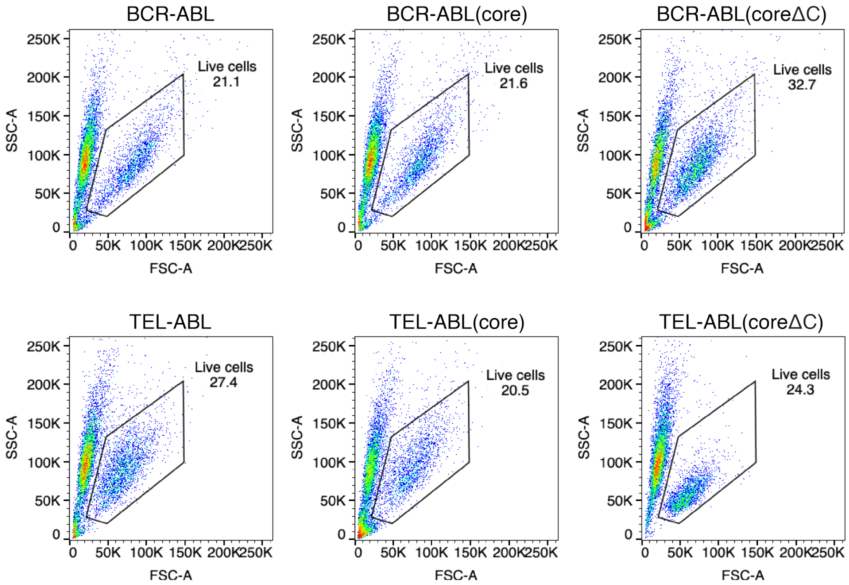


**Fig. S1.** Deleting the C-terminal segment of the ABL component does not affect the activity of either BCR-ABL or TEL-ABL. Live Ba/F3 cells transduced with the ABL fusion are gated in an FSC/SSC (forward scatter/side scatter) dot plot. All fusion constructs confer IL-3-independent growth.

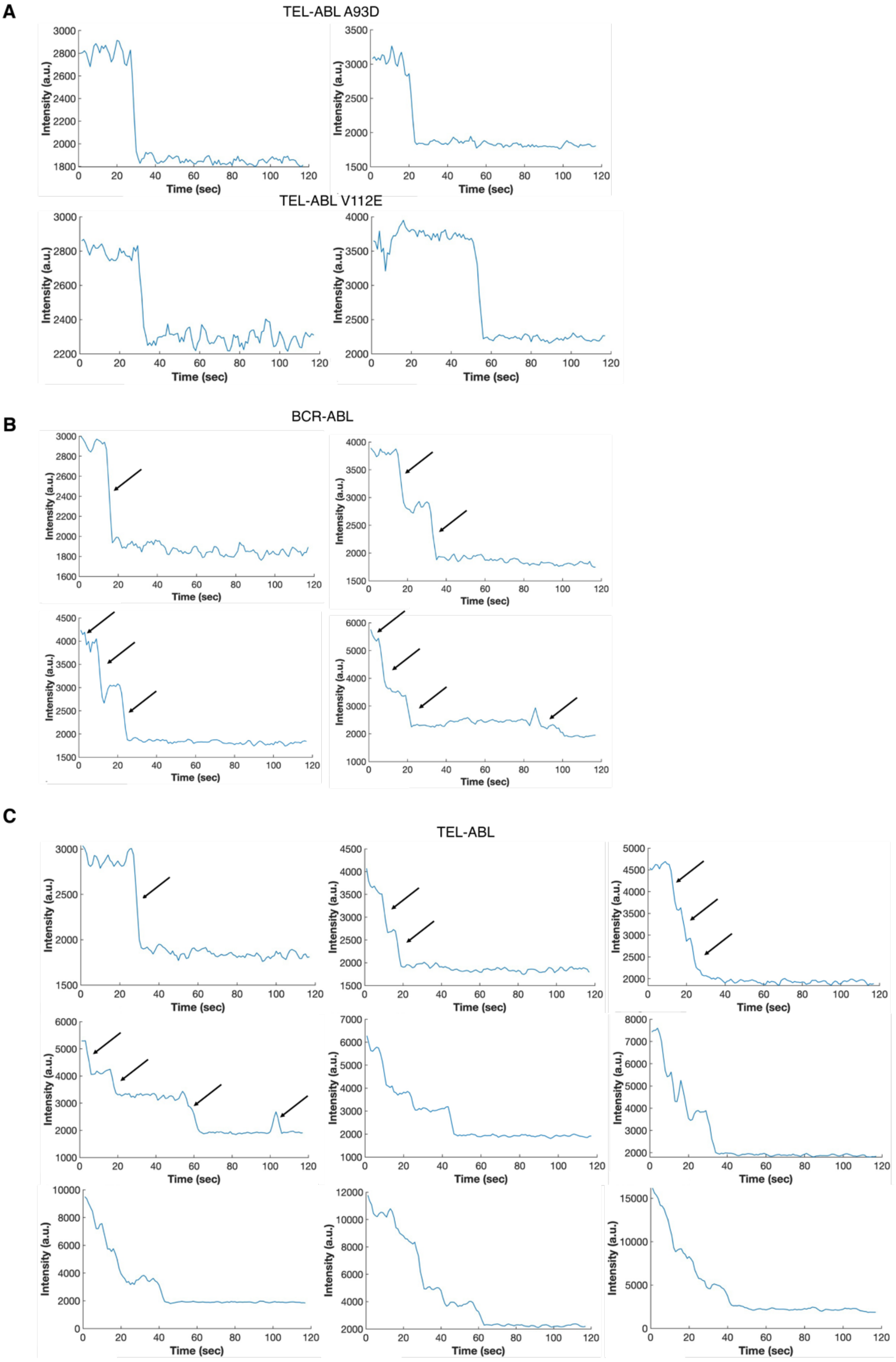


**Fig. S2.** Step photobleaching analyses of mNG-ABL fusion proteins. Intensity-versus-time records of **(A)** TEL-ABL A93D, TEL-ABL V112E, **(B)** BCR-ABL, and **(C)** TEL-ABL spots illustrating step photobleaching.

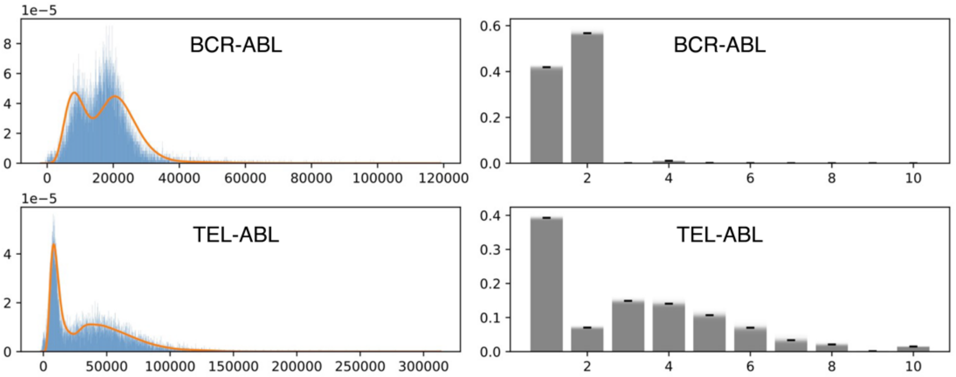


**Fig. S3.** Histogram of spot intensities for BCR-ABL (top) and TEL-ABL (bottom). The intensity probability density corresponding to the mean of parameter samples is overlaid in orange (left), and the distribution of oligomeric state distributions is shown in bar transparency, with the mean oligomeric state probability denoted with a horizontal line (right). Samples are from a model using the approximately-moment matched lognormal approximation to the sum of independent fluorophore intensities, and a fixed background intensity.

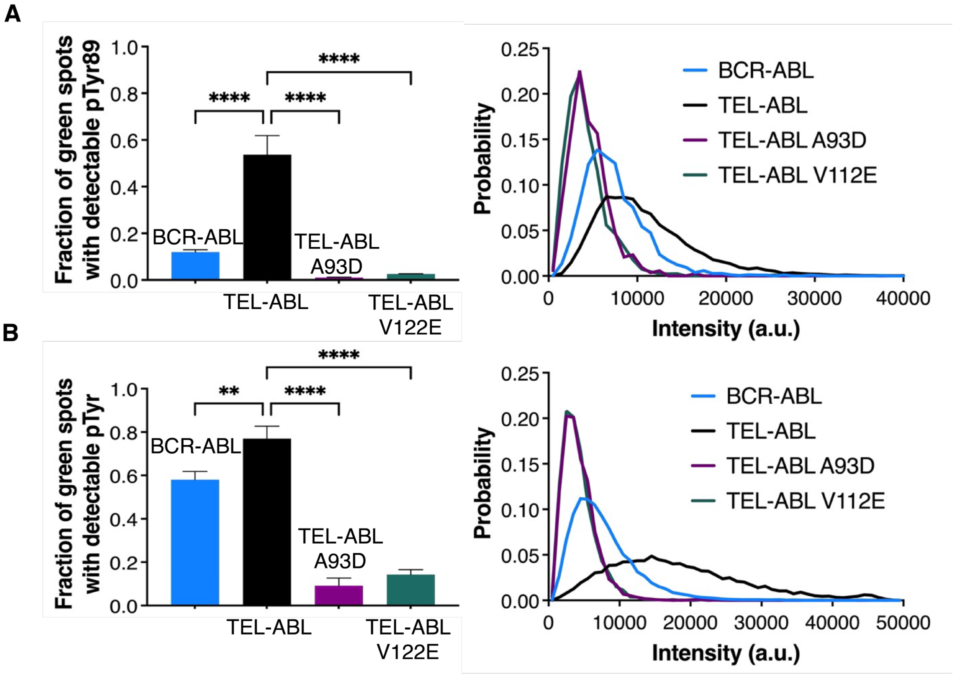


**Fig. S4.** The introduction of an ionizable residue into the TEL PNT domain reduces phosphorylation. **(A)** Fraction of BCR-ABL, TEL-ABL, TEL-ABL A93D, and TEL-ABL V112E that show detectable tyrosine 89 phosphorylation is plotted on the left. The distribution of intensities for pTyr 89 (561 nm) for mNG-ABL fusion proteins with detectable phosphorylation is plotted on the right (see Materials and Methods for details of normalization). **(B)** Fraction of BCR-ABL, TEL-ABL, TEL-ABL A93D, and TEL-ABL V112E that show detectable tyrosine phosphorylation is plotted on the left. The distribution of intensities for pTyr (561 nm) for mNG-ABL fusion proteins with detectable phosphorylation is plotted on the right (see Materials and Methods for details of normalization). (One-way ANOVA, N=3, number of experiments) (**** p≤0.0001, *** p≤0.001, ** p≤0.01, *p≤0.05, ns p>0.05).

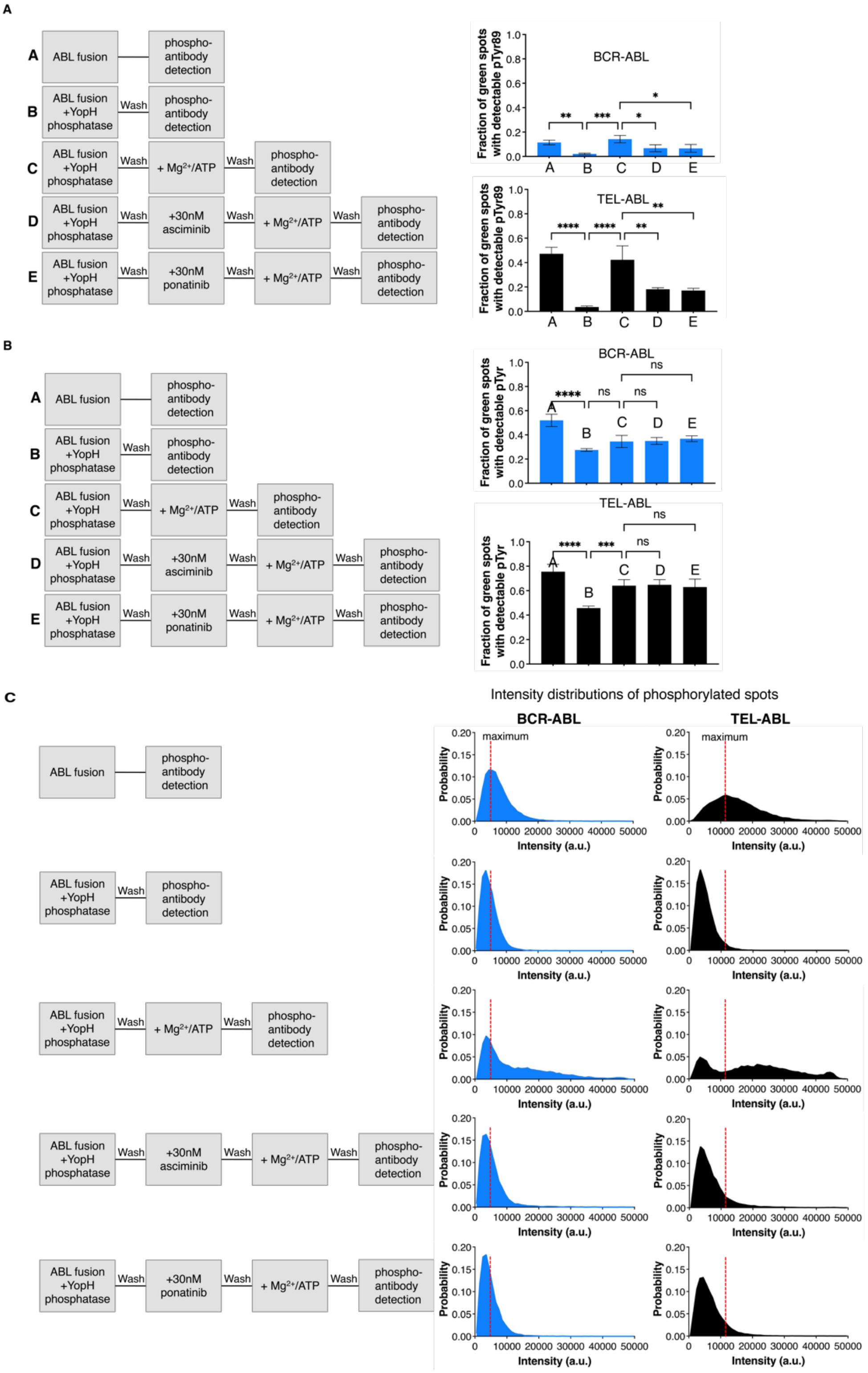


**Fig. S5.** TEL-ABL retains an intrinsic capacity to be inhibited by asciminib. **(A)** Fraction of BCR-ABL (in blue), and TEL-ABL (in black), at different reaction conditions, that show detectable tyrosine 89 phosphorylation is plotted. A schematic diagram showing the experimental setup is shown on the left. N=3 (number of experiments). **(B)** Fraction of BCR-ABL, and TEL-ABL, at different reaction conditions, that show detectable tyrosine phosphorylation is plotted. A schematic diagram showing the experimental setup is shown on the left. (One-way ANOVA, N=4, number of experiments). **(C)** The distribution of intensities for pTyr (561 nm), at different reaction conditions, for mNG-ABL fusion proteins with detectable Tyr phosphorylation is plotted on the right. A schematic diagram showing the experimental setup is shown on the left. N=4 (number of experiments, only the mean intensity distribution is plotted.) (**** $p \leq 0.0001$, *** $p \leq 0.001$, ** $p \leq 0.01$, *$p \leq 0.05$, ns $p > 0.05$).
